# Chromogranin A and catestatin regulate pancreatic islet homeostasis, endocrine function, and neurotransmitter signaling

**DOI:** 10.1101/2024.11.29.626063

**Authors:** Elke M. Muntjewerff, Dali Epremidze, Mariya Nezhyva, Satadeepa Kal, Theresa V. Rohm, Kechun Tang, Kailash Singh, Daniel Espes, Suborno Jati, Marleen Bootsma, Atef Mahmoud Mannaa, Hiromi Ikebuchi, Anna M. Nilsson, Mahadevan Rajasekaran, Per E. Andrén, Erik T. Jansson, Sushil K. Mahata, Gustaf Christoffersson

## Abstract

Chromogranin A (CgA), a neuroendocrine pro-hormone, undergoes proteolytic cleavage to yield bioactive peptides, notably catestatin (CST) and pancreastatin (PST), which exert opposing effects on metabolic and inflammatory processes. Using CgA and CST knockout (KO) mice, this study investigated their roles in pancreatic endocrine function, morphology, neurotransmitter dynamics, and systemic glucose homeostasis. CST deficiency induced insulin resistance, altered islet architecture, and heightened catecholamine levels, whereas CgA-KO mice lacking both CST and PST exhibited improved insulin sensitivity due to absence of PST. CST suppressed gluconeogenesis and enhanced glucagon regulation, while PST promoted insulin resistance and glucose production. Spatial mass spectrometry revealed altered neurotransmitter and polyamine profiles in pancreatic islets, implicating disrupted nerve-immune-islet interactions. CST’s modulation of catecholaminergic and inflammatory pathways positions it as a key regulator in the neuro-immune-endocrine axis. These findings highlight the therapeutic potential of targeting CgA-derived peptides, especially CST, for managing diabetes and metabolic-inflammatory diseases through precise peptide-based interventions.

## Introduction

Chromogranin A (CgA), a 49 kDa pro-hormone, has been extensively studied in the initiation and regulation of dense-core secretory granule biogenesis as well as in the context of various metabolic and inflammatory diseases. Increased circulating CgA levels are linked to various diseases including neuroendocrine tumors [1], hypertension [2,3] congestive heart failure [4], renal failure [5], inflammatory bowel disease [6], rheumatoid arthritis [7], sepsis [8], and Alzheimer’s disease [9,10]. CgA is secreted by endocrine cells such as chromaffin cells in the adrenal medulla, enterochromaffin cells in the gut, in pancreatic islets, neurons, and immune cells such as neutrophils, monocytes and macrophages [11]. In the pancreatic islet, CgA is present in the alpha and beta cells [12] and is important for islet volume, cellular composition and function [13]. CgA mostly exerts its effects following proteolytic cleavage, which is why most studies focus on the bioactive peptides. So far we know that CgA gives rise to vasostatin I/II (VS-I, VS-II), serpinin, WE-14, prochromacin, pancreastatin (PST) and catestatin (CST) [14,15]. The role of PST and CST have been investigated the most since they exert opposite effects [16]. For example, PST has anti-insulin and pro-inflammatory properties [17], while CST is pro-insulin and has anti-inflammatory properties [18,19].

Here, we mainly focus on the neuropeptide CST which exerts a wide variety of immune-and neuro-regulatory functions [11]. The 21 amino acid long CST acts as an anti-inflammatory [19–21], anti-diabetic [19] and cardioprotective peptide [22]. Decreased CST blood levels are seen in diet-induced obese mice [19] and type 2 diabetes (T2D) patients [23], while elevated CST blood levels are seen in inflammatory bowel disease [6], rheumatoid arthritis [24], COVID-19 infection [25] and various cardiovascular diseases such as acute heart failure, arrhythmia and hypertension [26–30]. Recent studies implicate that higher plasma CST, in combination with low plasma catecholamines, are associated with a worse disease prognosis for heart failure (five-fold increase in mortality for high plasma CST compared to normal CST) [26–30]. At the same time low plasma CST and high catecholamine levels are associated with insulin resistance [19], T2D [23] and other cardiovascular diseases including hypertension [3,23]. In line with this, the supplementation of CST in rodents with hypertension normalized blood pressure [21,31–33] with similar effects in humans [34]. CST supplementation also reduces chronic gut inflammation in murine colitis models [35,36] and improved inflammation and insulin sensitivity in mice with diet-induced obesity [19,37].

Based on the link between CgA and its cleavage products to various endocrine diseases and their presence in the pancreatic islets, we aimed to investigate how CgA, CST and PST may influence pancreatic islet endocrine function, islet morphology and microenvironment. For this purpose, we used CgA full knockout mice (CgA-KO) [31] and mice with selective deletion of the CST-coding region of the *Chga* gene (CST-KO) [21]. CgA-KO mice are characterized with hypertension, elevated sympathoneuronal activity and affected endocrine cells in the adrenal gland and pancreas [13,31]. CST-KO mice are hypertensive, obese and exhibit insulin resistance. Additionally, they display low grade organ inflammation, elevated amounts of norepinephrine and epinephrine in the plasma, lower bacterial gut diversity, and impaired epithelial barrier function [6,19,21]. CgA-KO mice maintained on a normal chow diet (NCD) exhibit enhanced insulin sensitivity [38], which has been attributed to the absence of the CgA cleavage product PST, a known inhibitor of hepatic gluconeogenesis [38]. Notably, insulin sensitivity in CgA-KO mice persists even after four weeks of high-fat diet [17], further reinforcing PST as a key contributor to insulin resistance. In contrast, CST-KO mice fed NCD display insulin resistance, suggesting that CST plays an insulin-sensitizing role. Both CgA-KO and CST-KO mice develop hypertension due to the absence of CST, as evidenced by the normalization of blood pressure upon CST supplementation [21,31]. Additionally, to dissect the opposing metabolic roles of PST (pro-diabetic) and CST (anti-diabetic), we supplemented CgA-KO mice with either peptide individually or in combination at equimolar concentrations to determine whether one peptide exerts dominant control over metabolic outcomes.

To our knowledge, this is the first study to comprehensively assess the pancreatic microenvironment, including endocrine cell density, composition and hormone content, pancreatic endocrine function by glucose-stimulated pancreatic hormone secretion and gluconeogenesis, inflammatory cytokine levels, innervation, and spatial mass spectrometry for identifying neurotransmitter and metabolite levels. We show that islets lacking CgA and CST display aberrant ratios of alpha- and beta cells along with glucagon and insulin contents and consequent differences in blood glucose homeostasis. Various neurotransmitters involved in crucial processes such as hormone regulation were also affected by CgA or CST deletion. In addition, we found that PST promoted gluconeogenesis, which was inhibited by CST and in combination, gluconeogenesis was restored to WT level. Our findings here thus make a major contribution in understanding the role of CgA and its cleavage products CST and PST in the pancreatic microenvironment with possible implications for the pathology of both T1D and T2D.

## 2 Results

### 2.1 CgA and CST in the pancreatic islet in homeostasis and in T1D

Since CST levels have been reported to be decreased [3,21,23] or increased in various inflammatory diseases [19,24–30], we measured plasma CST levels in healthy individuals and patients with T1D. In line with previous studies in patients with IBD [6] and RA [7], the CST levels were significantly increased in T1D individuals when compared to age-matched healthy controls (Fig. 1A). To visualize CST in the pancreas during autoimmune diabetes onset, we performed immunostainings in the pancreas of 4, 8 and 15 week-old female non-obese diabetic (NOD) mice. Quantification of the insulin and CST staining showed that the ratio of double-positive cells (insulin and CST) remained constant in the NOD mice islets indicating CST production throughout disease progress (Fig. 1B-C). At the moment, the exact CgA cleavage products present in the pancreas, including the ones leading to CST are unknown (Fig. 1D). To compare the presence of CgA cleavage products in wildtype and 8-week NOD male and female mice, we performed immunoblotting of pancreatic islets stained for CST (Fig. 1E, S-Fig. 1). To investigate tissue-specific processing of CgA into CST, we performed immunoblotting for CgA and CST in the adrenal medulla, pancreatic islets, and macrophages in WT mice (Fig. 1F, G, S-Fig. 2). Consistent with inhibition of proteolytic processing by catecholamines in the adrenal gland [39], we detected full-length CgA (75 kDa) in the adrenal medulla as compared to the pancreatic islets, which exhibited various CgA cleavage products, indicating rapid proteolytic processing of CgA (Fig. 1D, F). For both macrophages and pancreatic islets, CgA is effectively processed to CST as immunoblots with CST antibody showed bands comparable in size with the loaded synthetic CST peptide as shown previously in heart tissues [40] (Fig. 1F, G). Nevertheless, the presence of intermediate CgA cleavage products in the islets implies an active role for CgA peptides in maintaining islet homeostasis.

**Fig. 1.**
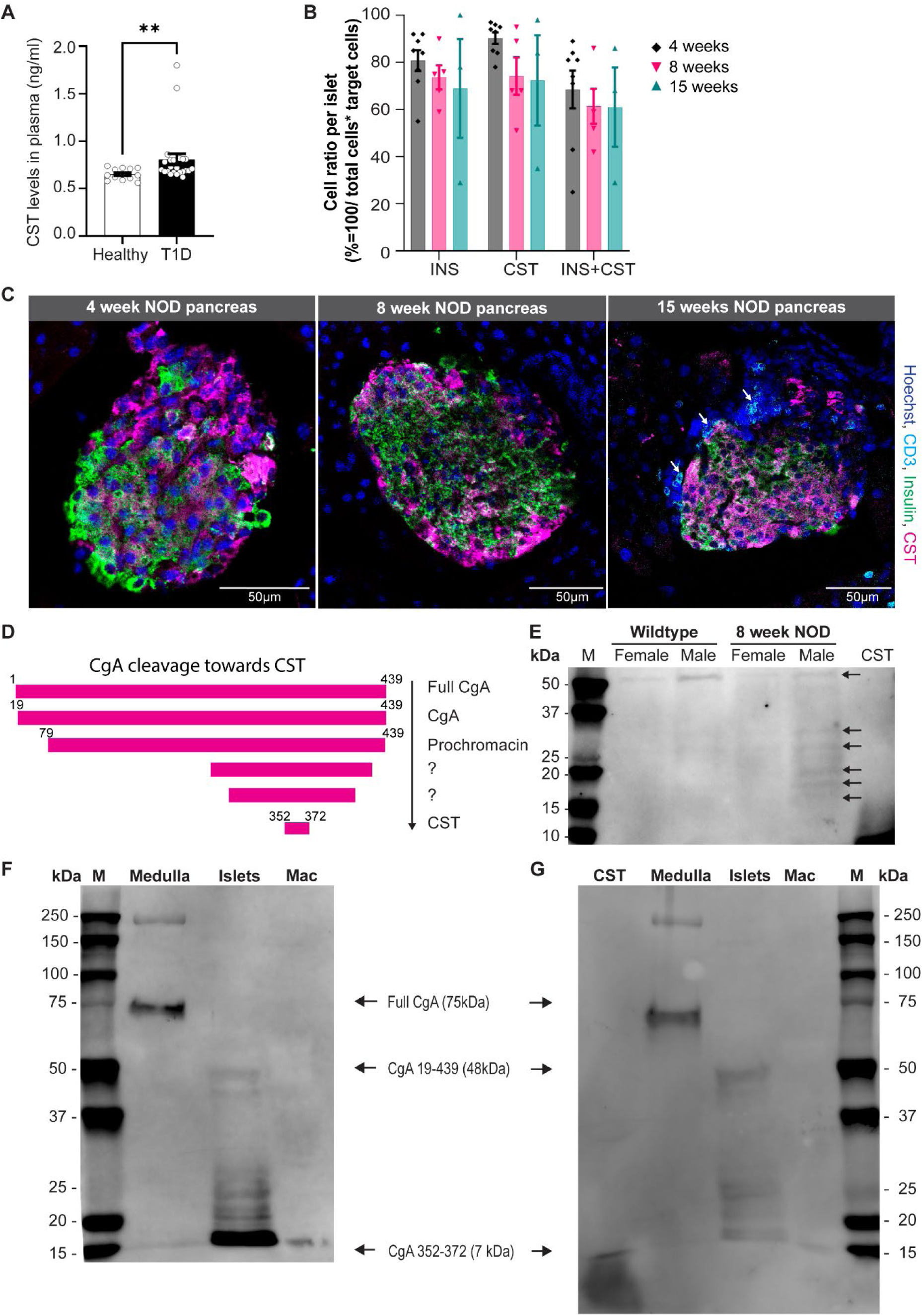
Chromogranin A (CgA) and catestatin (CST) in the pancreatic islet. **A)** CST plasma levels in healthy and T1D participants. **B-C)** Quantification and representative images of immunofluorescent staining for catestatin (CST, magenta), insulin (green), CD3 (cyan) and nuclei (blue) in 4, 8 and 15-week-old NOD mice pancreas **D)** CgA cleavage products towards CST. **E)** Western blot showing CST staining in NOD mice pancreas. **D)** Western blot showing CgA staining in the adrenal medulla, pancreatic islets and macrophages (mac). Arrows indicate full CgA and possible cleavage products of CgA. **E)** Similar membrane stained for CST with synthetic CST peptide in adrenal medulla, pancreatic islets, and mac.

### 2.2 Islet morphology and endocrine islet composition is altered upon CST or CgA deletion

Staining of frozen pancreas sections from WT, CgA-KO and CST-KO mice revealed CST staining in the pancreatic islets in WT mice. The complete absence of CST staining in the pancreata of CgA-KO or CST-KO knockout pancreata confirm their knockout phenotype (S-Fig. 3). We also observed CST staining outside the pancreatic islets in WT tissue, attributing to CST content in immune cells and/or nerve fibers. To get an overview of the pancreas of WT, CgA-KO and CST-KO sections, we made full-section scans of haematoxylin and eosin (H&E)-stained slides for male (Fig. 2A) and female mice (S-Fig. 5A). These images seemed to show changes in islet morphology, which could indicate aberrations in islet function [41,42]. To investigate pancreatic islet morphology and endocrine cell composition in CgA or CST-KO mice in more detail, we stained frozen pancreatic sections of KO mice for insulin, glucagon and somatostatin (Fig. 2B, S-Fig. 5B). First, we quantified the islet density and islet area. Although the individual islet areas were unchanged, the islet density was significantly decreased in the CgA-KO pancreas when compared to WT and CST-KO pancreas (Fig. 2C-D), which is in line with previous findings for CgA-KO mice [13]. Next, islet shape was assessed, showing decreased islet circularity in CST-KO compared to WT islets (Fig. 2E). To double check if the CST or CgA domain was absent in the pancreas, we ran a western blot with CST or CgA antibody, which showed no CST band for CST-KO mice and no CgA band in CgA-KO mice, confirming CST and CgA-KO (Fig. 2F, S-Fig. 4-6). To assess endocrine cell composition, we quantified alpha, beta and delta cell ratios per islet for WT, CgA-KO and CST-KO mice (Fig. 2F, S-Fig. 5C, S-Method Fig. 1). In line with previous findings [13], the quantification shows that CgA-KO mice display a significant increase in alpha cells and a decrease in beta cells per islet when compared to WT islets. The CST-KO mice only showed a decrease in beta cells when compared to the WT islets. In line with these findings, the numbers of hormone-negative cells in CST and CgA-KO islets were significantly increased. This means that CgA-KO and CST-KO mice have fewer endocrine cells per islet, which in combination with the decreased islet density in CgA-KO mice may lead to altered endocrine function. Thus, absence of CgA or CST in mice affects the endocrine cell composition and potentially also the function of the pancreatic islet.

**Fig. 2.**
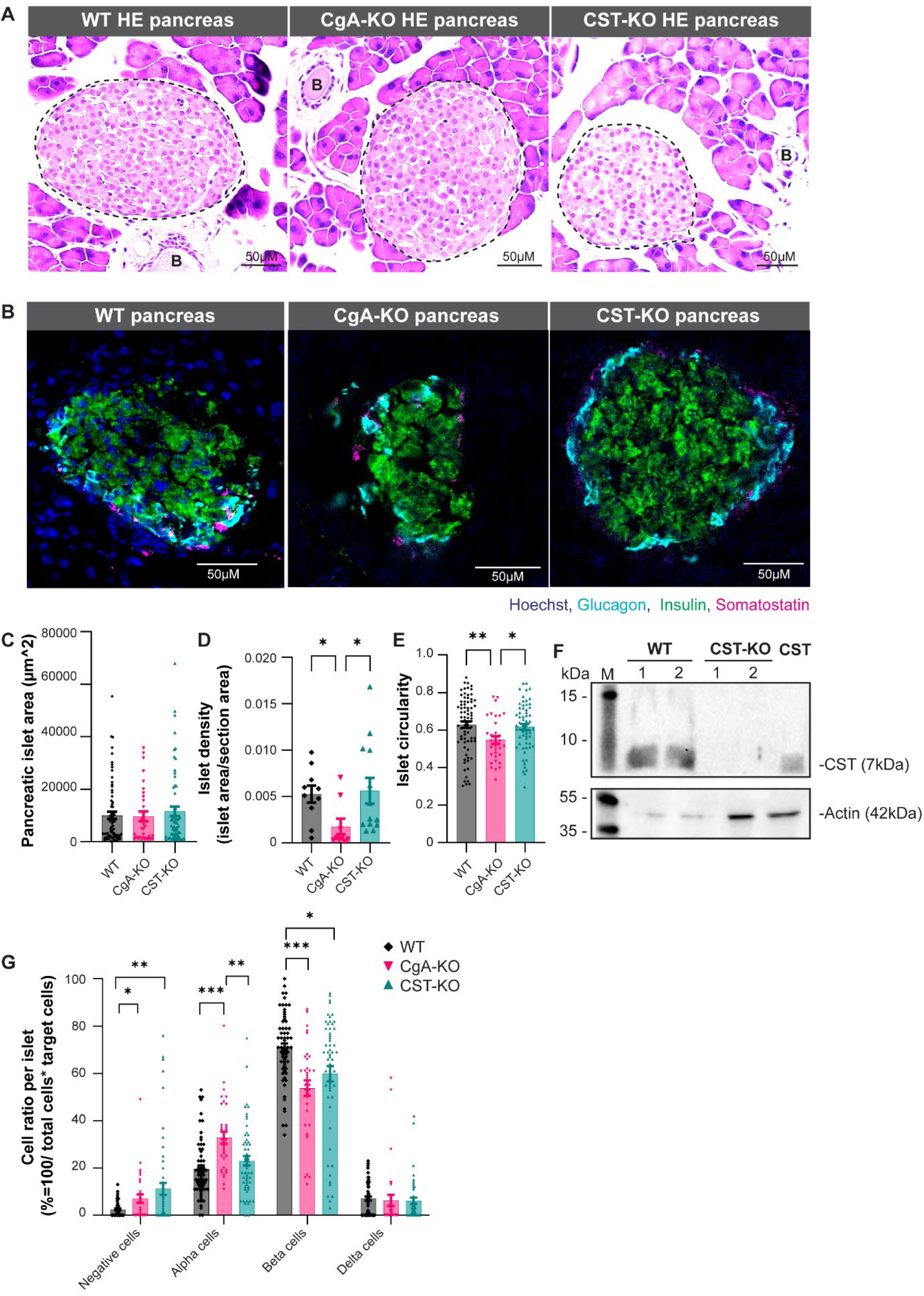
Composition of pancreatic islets in wild-type (WT), chromogranin A knockout (CgA-KO) and catestatin knockout (CST-KO) mice. **A)** H&E staining of WT, CgA-KO and CST-KO pancreatic slices including annotations for islets (white dotted lines) and blood vessels (B). **B)** Representative images of immunofluorescent stainings of glucagon (cyan), insulin (green), somatostatin (magenta) and Hoechst (blue) on WT, CgA-KO and CST-KO pancreatic sections. **C)** Islet area **D)** Islet density **E)** Islet circularity **F)** Immunoblotting showing CST staining in pancreatic islets of WT and CST-KO mice with synthetic CST peptide (CST) as a positive control. **G)** Quantification of alpha/beta/delta/hormone negative cells per islet. Quantification is based on the staining represented in panel B. n=5 mice per group. *p<0.05 , **p<0.01, ***p<0.001

### 2.3 Pancreatic islet function of mice lacking CgA or CST

To determine the functional consequences of CgA or CST deficiency on glucose homeostasis and islet endocrine response, we performed *in vivo* glucose tolerance tests (GTT) in male mice. CgA-KO mice exhibited significant reduction (∼17% lower AUC) in blood glucose levels during GTT compared to WT controls (Fig. 3A, B). This is consistent with the enhanced peripheral insulin sensitivity, likely due to suppression of hepatic gluconeogenesis, as reported previously [38]. These physiological findings support earlier reports of reduced islet number, decreased β-cell mass, and lower plasma insulin levels in CgA-KO mice relative to WT [13]. Conversely, CST-KO mice displayed significant elevation (∼18% higher AUC) of blood glucose levels during GTT compared to WT mice, indicative of impaired β-cell function and/or insulin resistance, as described previously [19].

**Fig. 3.**
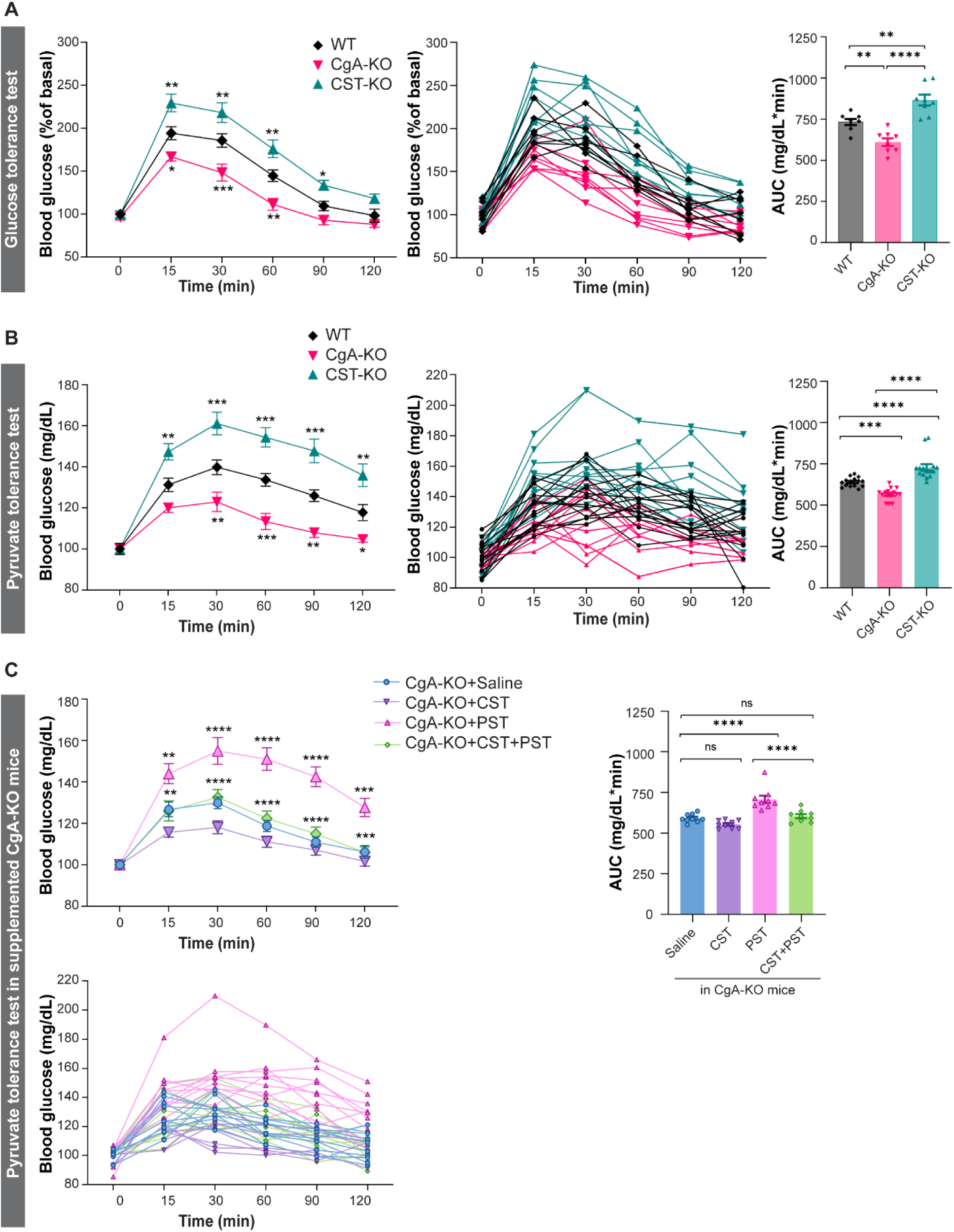
Pancreatic islet function of mice lacking chromogranin A (CgA) or catestatin (CST) **A)** Graphs displaying combined and single data of glucose tolerance test (GTT; 2g/kg dextrose intraperitoneal) results for WT (black), CgA-KO (pink) and CST-KO (green) over time (min) n=8 mice per group (Statistical Power = 95%) including a graph displaying the calculated area under the curve (AUC in mg/dL*min). **B)** Graphs displaying combined and single data of pyruvate tolerance test (PTT: i.p. 2 g/kg bw sodium pyruvate) blood glucose results (mg/dL) for WT (black), CgA-KO (pink) and CST-KO (green) over time (min) n=15 mice per group (Statistical Power >95%) and a graph displaying the calculated AUC **C)** Graphs displaying combined and single data of PTT blood glucose results (mg/dL) over time (min) of CgA-KO mice supplemented with saline, CST, PST or CST+PST. n=10 mice per group (Statistical Power = 95%) and a graph displaying the calculated AUC. Data were analyzed by one- or two-way ANOVA followed by Sidak’s multiple comparison test. *p<0.05, **p<0.01, ***p<0.001, ****p<0.0001

Given the central role of gluconeogenesis in glucose homeostasis and its suppression by insulin, we next conducted pyruvate tolerance tests (PTT) in WT, CgA-KO, and CST-KO mice to assess gluconeogenic capacity. We recorded body weight, pancreatic weight and food intake before starting PTT in male and female WT, CgA-KO and CST-KO mice (S-Fig. 7-8), which showed increased body weight for CgA-KO and CST-KO male and female mice compared to WT mice. The CST-KO mice did not only increase in weight, but also lost the normal difference in weight between male and female mice. To access how efficiently these mice convert pyruvate into glucose via gluconeogenesis, we conducted PTT. Consistent with a previous study [38], CgA-KO mice exhibited marked reduction (∼11% and ∼8% lower AUC in male and female mice, respectively) in glucose levels during PTT (Fig. 3C, S-Fig. 9-10), indicating inhibition of hepatic gluconeogenesis. In contrast, CST-KO mice showed increased (∼14% and ∼13% higher AUC in male and female mice, respectively) glucose levels (Fig. 3C, S-Fig. 10A-D), suggesting enhanced gluconeogenesis in the absence of CST. In summary, deletion of CgA or CST results in divergent glucose metabolism. Deletion of CgA, results in heightened insulin sensitivity, while deletion of only the CST domain results in decreased insulin sensitivity.

To delineate the contribution of specific CgA-derived peptides to the regulation of gluconeogenesis, we supplemented CgA-KO mice intraperitoneally with PST (1.4 µg/g body weight), CST (1 µg/g body weight), or an equimolar concentration (215 µM) of both and performed PTT after four weeks of treatment. We recorded body weight throughout the treatment and pancreas weight was recorded after sacrificing mice for tissue harvesting. Male and female CgA-KO mice supplemented with saline, CST and CST+PST showed a similar pattern as for the normal CgA-KO mice (S-Fig. 7-10). CST-treated CgA-KO mice showed decreased glucose production (∼7% and ∼11% decrease in AUC in male and female mice, respectively) (Fig. 3D, S-Fig. 10E), indicating inhibition of gluconeogenesis by CST specifically in female mice. In contrast, PST-treated mice exhibited increased (∼19% and ∼22% increase in AUC in male and female mice, respectively) glucose levels (Fig. 3D, S-Fig. 10E), implicating PST as a pro-gluconeogenic peptide. Notably, co-administration of CST and PST prevented the PST-induced rise in glucose levels during PTT (Fig. 3D, S-Fig. 10E), indicating that CST functionally antagonizes PST’s effects and exerts dominant control over gluconeogenesis. Thus, it seems that PST and CST have opposite effects in gluconeogenesis, where PST promotes gluconeogenesis and CST suppresses gluconeogenesis. The above findings are consistent with heightened insulin sensitivity in CgA-KO mice [38] owing to the lack of PST and increased insulin resistance in CST-KO mice [19] owing to the lack of CST.

### 2.4 Glucose-stimulated insulin secretion from isolated CgA-KO and CST-KO pancreatic islets

To assess islet functionality further, we performed glucose-stimulated insulin secretion (GSIS) assays on islets isolated from WT, CgA-KO and CST-KO mice. The islet yields per animal tended to be slightly higher for CST-KO mice when compared to WT and CgA-KO (Fig. 4A). Total insulin content per 10 islets was comparable across all genotypes, but insulin levels were slightly increased in the CST-KO islets (Fig. 4B). High glucose-stimulated insulin secretion was comparable between WT and CgA-KO mice (∼12-fold versus ∼13-fold) as compared to low glucose (Fig. 4C). In contrast, in CST-KO mice, high glucose caused ∼29-fold increase in insulin secretion (>2-fold higher than CgA-KO islets Fig. 4C). The fold changes in insulin secretion in response to high glucose were as follows: ∼10-fold in WT islets; ∼13-fold in CgA-KO islets and ∼33-fold in CST-KO islets (Fig. 4D). Insulin secretion index was as follows: ∼10-fold for WT islets; ∼14-fold for CgA-KO islets and ∼36-fold for CST-KO islets. These data show a major increase in the *ex vivo* response to glucose from CST-KO islets compared to the *in vivo* GTT data.

**Fig. 4.**
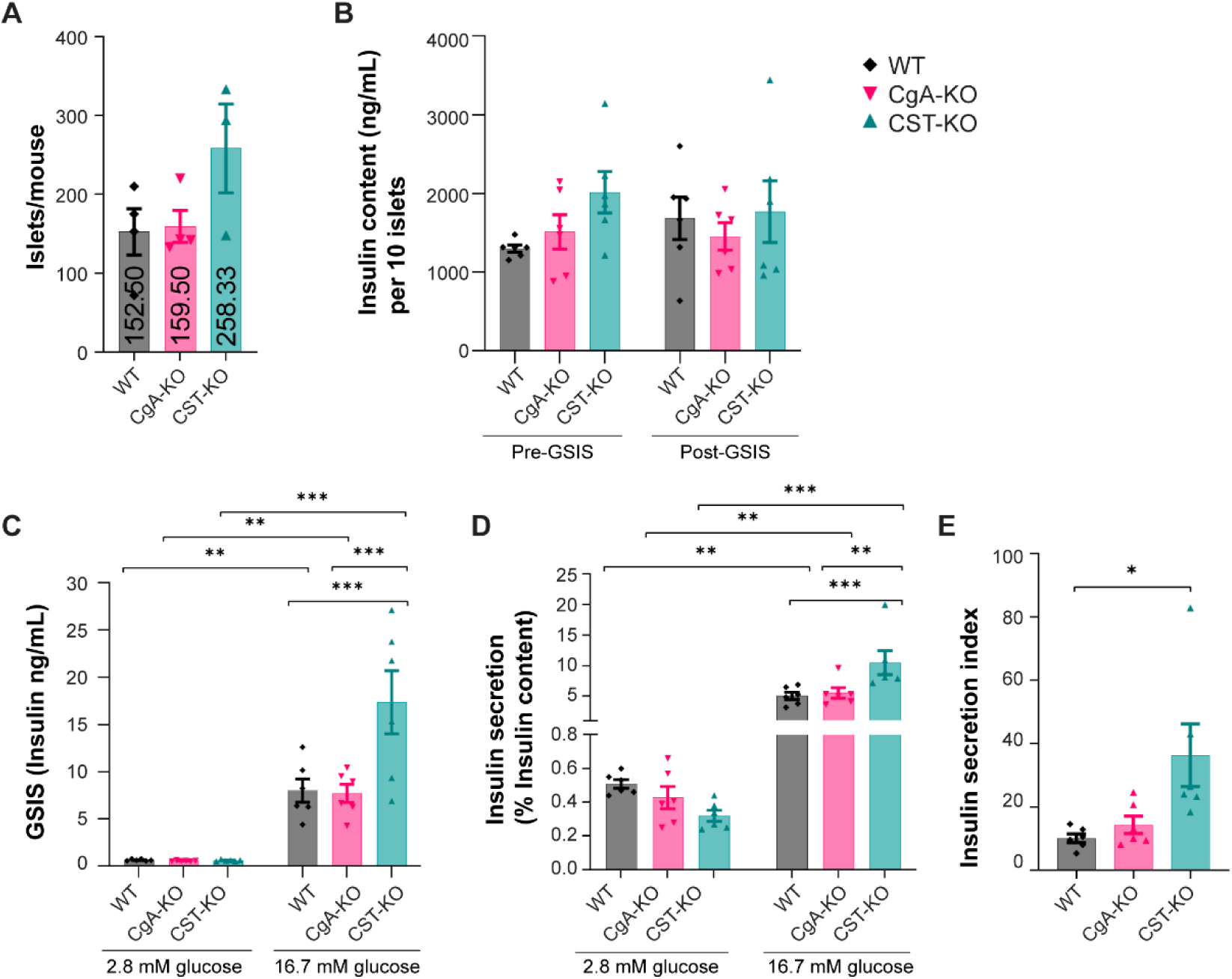
Glucose-stimulated insulin secretion (GSIS) from islets of mice lacking chromogranin A (CgA) or catestatin (CST) **A)** Islet yield per animal following isolation from WT, CgA-KO and CST-KO pancreata. **B)** Total insulin content (ng/mL) per 10 islets before (pre) and after (post) glucose stimulation for WT, CgA-KO and CST-KO mice **C)** GSIS from isolated islets from WT, CgA-KO and CST-KO mice. Showing insulin content under basal glucose conditions (2.8 mM) before glucose stimulation (pre-GSIS) (left) and after high glucose stimulation (16.7 mM) (post-GSIS) (right). **D)** Insulin secretion during GSIS on isolated islets from WT, CgA-KO and CST-KO mice, expressed as percent of insulin content before and after GSIS. **E)** Insulin secretion index = fold change of high vs. low GSIS (/content). n=6 mice per group (Statistical Power = 80%). Data were analyzed by one- or two-way ANOVA followed by Sidak’s multiple comparison test. *p<0.05, **p<0.01, ***p<0.001, ****p<0.0001

### 2.5 *In vivo* insulin and glucagon secretion after supplementation of CgA-KO mice with CST, PST or CST+PST

To investigate the role of CgA in islet endocrine function, we first assessed fasting (8 h) plasma insulin, glucagon, and C-peptide levels in male WT, CgA-KO, and CST-KO mice. Compared to WT mice, fasting plasma insulin levels were significantly reduced in CgA-KO (∼28%) and CST-KO (∼54%) mice (Fig. 5A). Decreased plasma insulin in CST-KO mice coincides with decreased C-peptide level (S-Fig. 7A). Paradoxically, fasting plasma glucagon levels were significantly decreased only in CST-KO mice (∼50%), but not in CgA-KO mice (Fig. 5B). To further elucidate islet secretory dynamics, we conducted GTTs. Glucose administration in fasting (8 h) mice caused ∼1.3- and ∼1.6-fold increase in insulin secretion in WT and CgA-KO mice, respectively as compared to only ∼1.1-fold in CST-KO mice (Fig. 5C), in line with the increased glucose levels (Fig. 3A and B). Suppression of glucagon secretion by glucose was observed only in fasting WT mice (∼40%), but this response was abolished in both CgA-KO and CST-KO mice (Fig. 5D), implicating CST as a key regulator of islet glucagon dynamics.

**Fig. 5.**
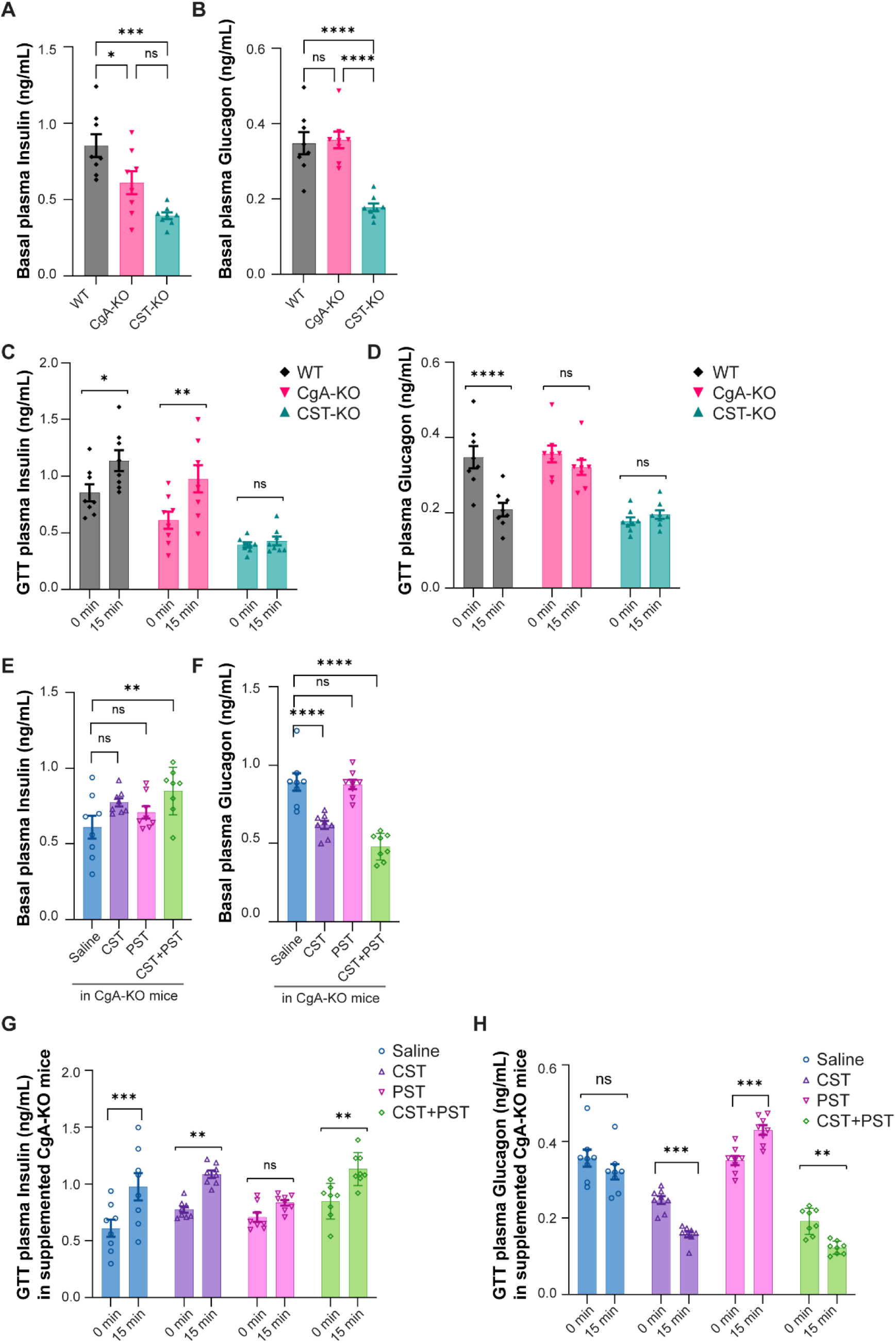
Basal and glucose stimulated hormone release upon supplementation or lack of chromogranin A (CgA)-cleavage products. **A)** Basal fasting plasma insulin and **B)** glucagon levels (ng/mL) from WT, CgA-KO and CST-KO mice. **C)** Graph showing plasma insulin and **D)** glucagon concentrations under basal glucose conditions (fasting) and under high glucose conditions (15 min after i.p. injection of 2 g/kg dextrose) for WT, CgA-KO and CST-KO mice, n=8 mice per group **E)** Basal fasting plasma insulin and **F)** glucagon levels (ng/mL) in CgA-KO mice treated for seven days with saline, CST, PST or CST+PST. **G)** Graph showing fasting plasma insulin or **H)** glucagon levels under basal glucose conditions and under high glucose conditions (15 min after i.p injection of 2 g/kg dextrose) injected for 4 weeks with saline, CST (1 µg/g bw, i.p.), PST (1.4 µg/g bw, i.p.) or CST+PST. Data were analyzed by one- or two-way ANOVA followed by Sidak’s multiple comparison test. n=8 mice per group (statistical power = 85%). *p<0.05, **p<0.01, ***p<0.001, ****p<0.0001

To dissect the contributions of CgA-derived peptides associated with peripheral insulin resistance, we supplemented CgA-KO mice with intraperitoneal PST, CST, or a combination of both peptides at equimolar concentrations for one month. Basal plasma insulin levels were not affected significantly after treatments with CST or PST (Fig. 5E). However, co-administration of CST and PST significantly elevated (by ∼39%) fasting insulin levels (Fig. 5E), suggesting that CST counteracts the effects of PST and enhanced basal insulin secretion. With respect to glucagon, CST monotherapy significantly reduced (by ∼31%) fasting plasma glucagon levels (Fig. 5F), while PST had no effect. Notably, combined CST and PST treatment resulted in further suppression of glucagon (by ∼46%; Fig. 5F), highlighting CST’s dominant role in regulating glucagon secretion.

GTT experiments in peptide-treated CgA-KO mice revealed: (i) enhanced glucose-induced insulin secretion (∼60%) in saline-treated controls; (ii) moderate insulin secretion (∼40%) with CST treatment; (iii) inhibition of insulin secretion by PST; and (iv) restoration of glucose-stimulated insulin secretion (∼33%) in the presence of both CST and PST (Fig. 5G), indicating that CST is able to override PST-mediated inhibition. Finally, while glucose failed to suppress glucagon secretion in saline-treated CgA-KO mice, CST or CST+PST treatment restored this physiological suppression (∼36%; Fig. 5H). In contrast, PST treatment alone led to paradoxical stimulation of glucagon secretion in response to glucose (Fig. 5H).

### 2.6 Neurotransmitter and metabolite levels are changed in CgA-KO and CST-KO pancreas

Neuronal input to islet endocrine cells affects their hormonal secretion. In order to explore potential differences in pancreatic neuronal activity, we employed mass spectrometry imaging of pancreatic sections from WT, CgA-KO and CST-KO. We detected 12 pancreatic neurotransmitters and metabolites that met validation criteria (S-Table 2). Values were extracted for WT, CgA-KO and CST-KO total pancreas and differential expressions were visualized in a bar chart (Fig. 6A, S-Fig. 11-12) These data showed that hypoxanthine, tyrosine, spermidine, histidine, cysteine, spermine levels differ between CST-KO and WT. For most neurotransmitters, the expression seemed to be lowest in the CgA-KO pancreas overall when compared to WT and CST-KO. To be able to distinguish neurotransmitter levels between pancreatic islets and exocrine pancreatic tissue, we identified islets on consecutive sections stained with H&E (S-Fig. 13). After scanning the pancreatic slices in a microscope, we matched the exported islet regions to the intensity distribution of the serotonin signal and annotated islets and comparatively sized areas of exocrine tissue (serotonin negative) (Fig. 6B, S-Method Fig. 2-3). Here, we visualized and quantified the content of serotonin, GABA, histamine, cysteine, taurine, creatine, spermidine, histidine, norepinephrine (NE), tyrosine, spermine and hypoxanthine in the exocrine and endocrine parts of the pancreas (Fig. 6C). Altogether this resulted in a comprehensive view of the neurotransmitters in the WT, CgA-KO and CST-KO pancreas.

**Fig. 6.**
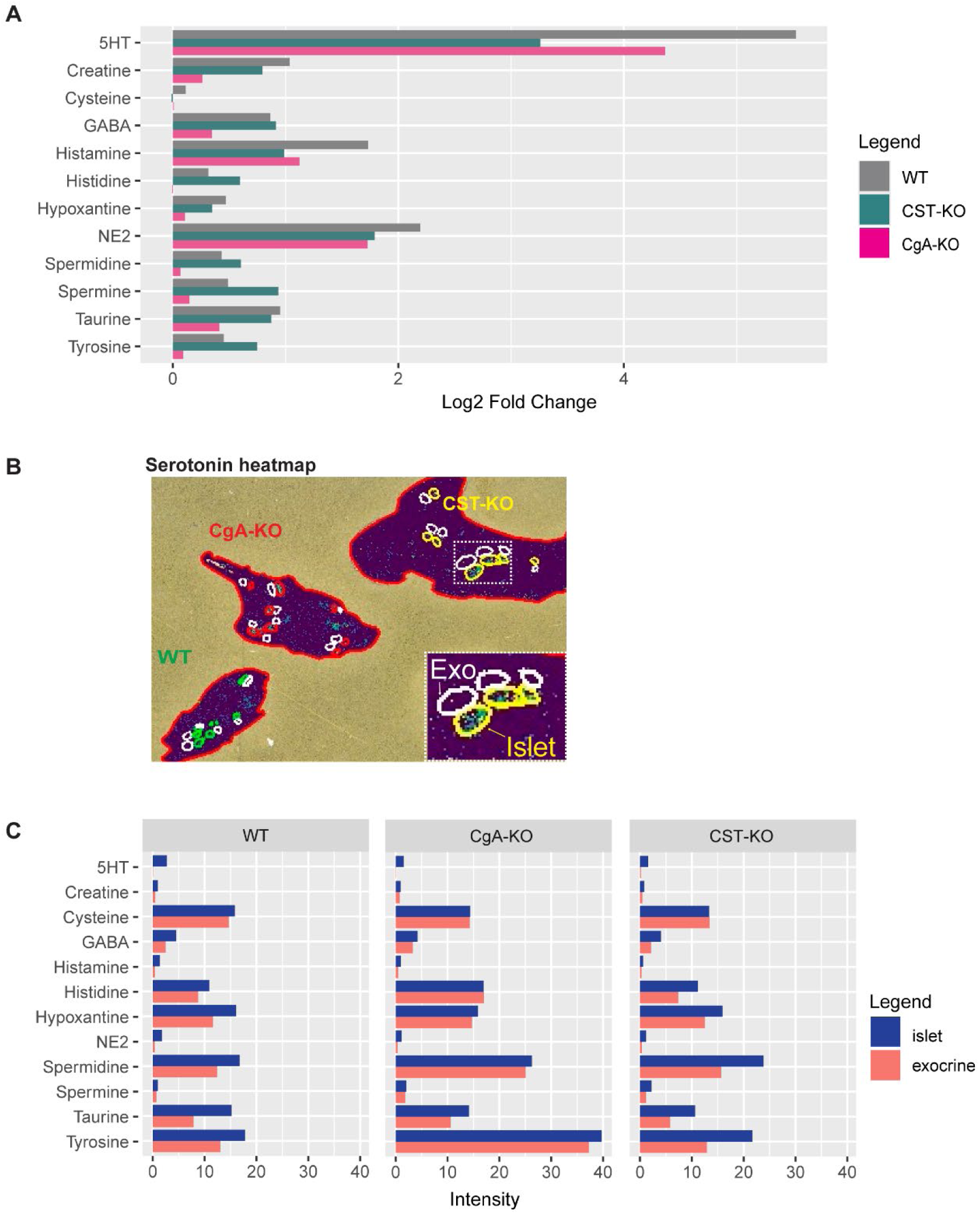
Pancreatic neurotransmitter regulation upon chromogranin A (CgA) or catestatin (CST) knockout. **A)** Comparative analysis of analytes in pancreatic tissue. This chart displays the log_2_-fold change in expression of the analytes in total pancreatic tissue of WT (gray), CgA-KO (pink) and CST-KO (blue). **B)** Serotonin heatmap showing examples of WT (green annotations), CgA-KO (red annotations) and CST-KO (yellow annotations) pancreatic sections including annotations for islets (colored) and exocrine tissue (white). **C)** Comparative analysis of analytes in pancreatic tissues. This figure highlights the differential expression of neurotransmitters between pancreatic islets (blue) and adjacent exocrine tissue (red), based on the average intensity of root mean square (RMS) normalized regions. n= 3 mice per group.

### 2.7 Metabolic and neurotransmitter synthesis pathways are affected in CgA-KO and CST-KO pancreatic islets

In addition to the combined analysis, we took a closer look at the individual analytes in pancreatic islets and exocrine pancreas to identify which parts of the pathways were affected in islets and exocrine tissue of CgA-KO and CST-KO pancreata. For all three genotypes, serotonin and taurine levels were significantly higher in pancreatic islets when compared to exocrine tissue (Fig. 7A-B). Additionally, GABA and histamine levels were higher in islets when compared to exocrine tissue for WT and CST-KO (Fig. 7D-E). We also noted that CgA-KO had lower GABA levels in islets, and for both GABA and histamine no differences between islet and exocrine tissue were detected. For CgA-KO islets and exocrine tissue we also observed a decrease in amounts of cysteine compared to WT (Fig. 7F). Cysteine is a highly conserved amino acid involved in regulating catalysis, protein structure, redox sensitivity and metal-ion transport [43]. Decreased cysteine levels might indicate disturbances within the homocysteine pathway.

**Fig. 7.**
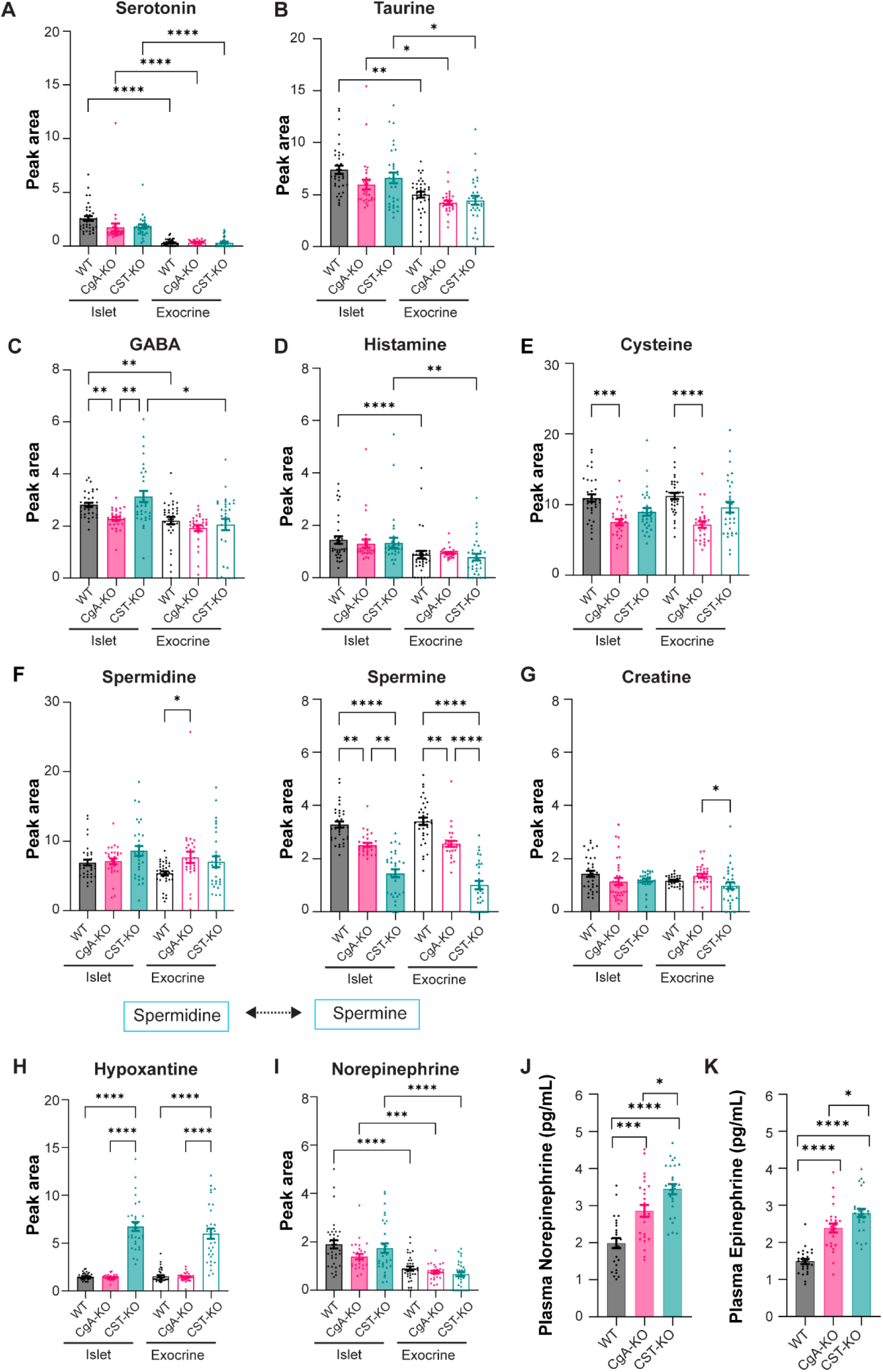
Islet and exocrine pancreatic neurotransmitter regulation upon chromogranin A (CgA) or catestatin (CST) knockout. **A)** Peak area intensities for islets and exocrine tissue of WT, CgA-KO and CST-KO pancreata of serotonin, **B)** taurine **C)** GABA **D)** histamine **E)** cysteine **F)** spermidine and spermine. Spermidine can be converted into spermine as illustrated in the schematic drawing. **G)** Peak area intensities of creatine for islets and exocrine tissue of WT, CgA-KO and CST-KO pancreata **H)** hypoxantine **I)** norepinephrine. n= 3 mice per group. **J)** Plasma norepinephrine (pg/mL) and **(K)** epinephrine (pg/mL) of WT, CgA-KO and CST-KO mice, n=26 mice per group (Statistical Power = >95%). Data were analyzed by one-way ANOVA followed by Sidak’s multiple comparison test. *p<0.05, **p<0.01, ***p<0.001, ****p<0.0001

Polyamines (spermidine, spermine and thermospine) play important roles in cell growth, proliferation and differentiation. In the pancreas, polyamines seem to modulate beta cell function by affecting proinsulin biosynthesis and insulin secretion [44]. Spermidine is synthesized from putrescine by spermidine synthase. This process takes place in all mice since similar spermidine values were found for WT, CST-KO and CgA-KO pancreas (Fig. 7G). Afterwards, spermidine is normally converted into spermine by spermine synthase. However, this process seems disturbed in both knockout models as spermine levels were drastically decreased for the CgA-KO and CST-KO islets and exocrine tissue when compared to WT pancreas (Fig. 7G). This suggests that the conversion by spermine synthase is less efficient in the absence of CgA or CST.

The creatine pathway seems unaffected since creatine levels appear equal for all three genotypes (Fig. 7G). Hypoxanthine levels are only significantly increased in CST-KO mice (Fig. 7H). Here both exocrine tissue and islets showed high levels of hypoxanthine, which could indicate pancreatic necrosis [45].

Norepinephrine (NE) and dopamine are both catecholamines that can regulate insulin secretion by islet beta cells [44,46]. Thereby the synthesis and degradation of NE and dopamine are essential for normal islet endocrine function. In the past, increased adrenal and plasma catecholamine levels have been detected in the plasma and adrenal gland of CgA-KO and CST-KO mice [21]. In contrast, our data shows that NE levels in the pancreas are not significantly different compared to WT for CgA-KO and CST-KO mice since we observed high islet NE levels and lower NE exocrine levels for all three genotypes (Fig. 7I). These data suggest that catecholamine levels in the pancreas were normal. In contrast, catecholamine plasma levels (NE and epinephrine), were increased in CgA or CST deletion (Fig. 7J, K), indicating that there might be a disturbance in local catecholamine degradation or an influence from circulating catecholamines.

### 2.8 Pancreatic sympathetic innervation and inflammatory signature in mice lacking CgA and CST

There is increasing evidence that pancreatic islet innervation is disordered in T1D and T2D, which may lead to changes in regulation of islet hormone release and metabolic regulation [47]. To investigate whether the observed changes in endocrine composition and/or function in CgA-KO and CST-KO mice may be related to islet innervation, we stained pancreatic sections for the norepinephrine transporter (NET) to visualize sympathetic neurons (Fig. 8A, S-Fig. 14A). Although the nerve network qualitatively seemed more disorganized in the CgA-KO and CST-KO pancreas, the quantification did not reveal major measurable differences regarding islet innervation between WT, CgA-KO and CST-KO pancreata (Fig. 8B). Comparing innervation in and around the pancreatic islet, the nerve density for WT islets was significantly lower when compared to the exocrine nerve network (Fig. 8C). However, this difference between in- and outside the islet was lost for the CgA-KO and CST-KO mice. This might indicate changes in the overall nerve network organization upon CgA or CST deletion. CST acts as a neuropeptide and has been shown to be produced by and have effect on nerves and macrophages, which may result in suppression of neuronal and neuroendocrine activity in an inflammation-dependent manner [11]. To investigate its role in the pancreas, we quantified nerve-macrophage interactions in pancreatic islets of CgA and CST-KO mice (Fig. 8D-F). Although no difference was found in the number of macrophages per islet, we observed a trend for less macrophage-nerve interactions in the islet upon CgA or CST deletion when compared to WT (Fig. 8E, F). To gain a better insight into the role of inflammation in mice lacking CgA or CST, we measured plasma TNF-α, interleukin-6 (IL-6), IL-1β, C-X-C motif chemokine ligand 1 (CXCL-1), chemokine (C-C motif) ligand 2 (CCL2), and IL-10. In line with previous studies [6,19,21], the lack of CST resulted in reduced plasma levels of the anti-inflammatory marker IL-10 and increased inflammatory cytokines (IL-6, IL-1β) and chemokine levels (CXCL-1, CCL-2), reinforcing CST as an anti-inflammatory peptide (Fig 8G).

**Fig. 8.**
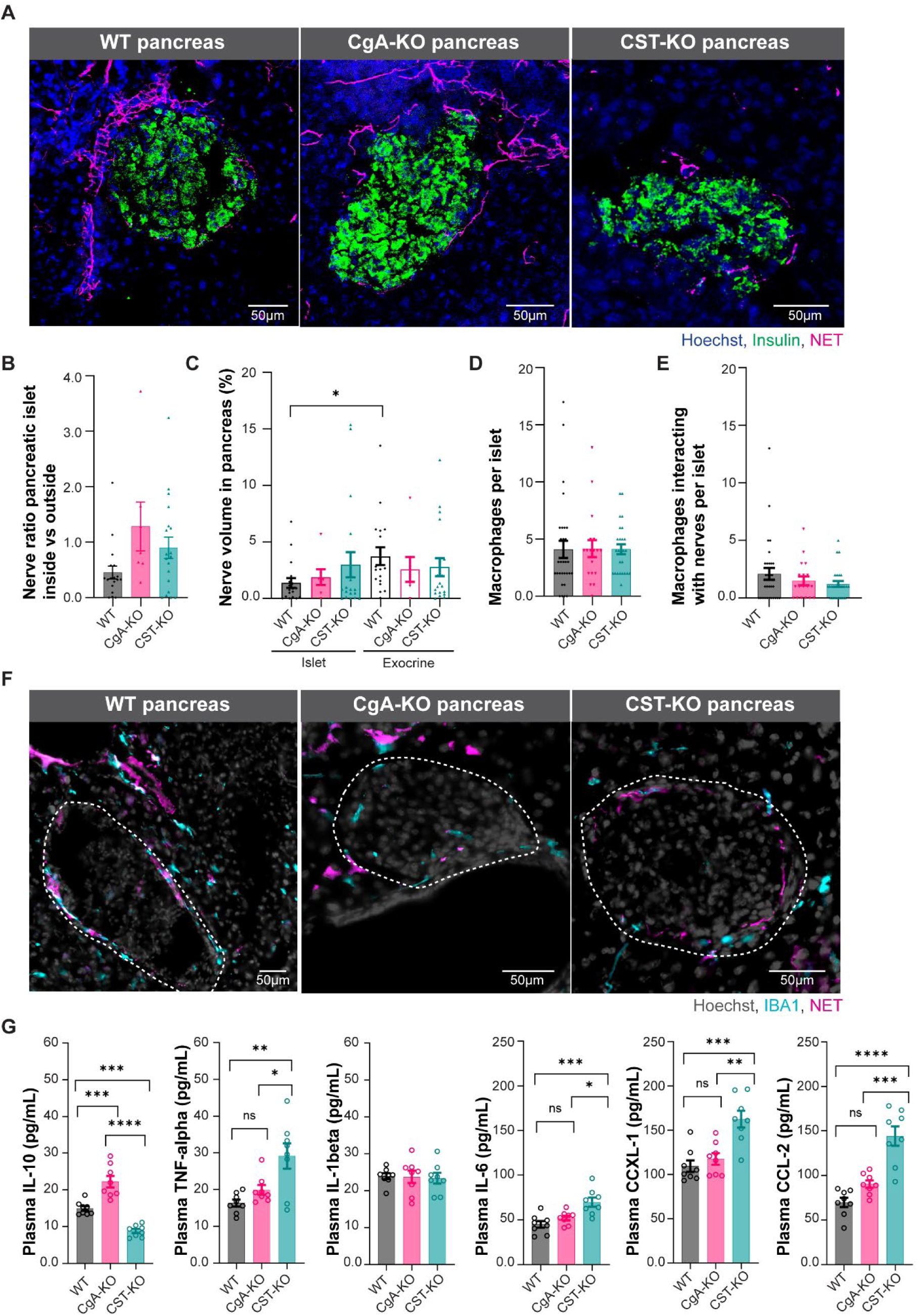
Innervation of pancreatic islets and plasma cytokines in wild-type (WT), chromogranin A knockout (CgA-KO) and catestatin knockout (CST-KO) mice. **A)** Representative images of nerve staining (norepinephrine transporter (NET, magenta), islets (insulin, green) and nuclei (Hoechst) in WT, CgA-KO and CST-KO pancreatic sections. **B)** Quantification of the nerve density ratio between islet and exocrine tissue for WT, CgA-KO and CST-KO based on the stainings represented in (A). **C)** Quantification of nerve volume in the islet and the exocrine pancreas for WT, CgA-KO and CST-KO. **D)** Macrophage count per islet (every dot in the chart represents one islet). **E)** Number of macrophages interacting with nerves per islet for WT, CgA-KO and CST-KO. N=3 mice per group. **F)** Representative images of nerve staining (norepinephrine transporter (NET, cyan), macrophages (IBA1, magenta), islets (insulin, green) and nuclei (gray) in WT, CgA-KO and CST-KO pancreatic slices. **G)** Plasma IL-10, TNF-α, IL-β, IL-6, CXCL-1 and CCL-2 levels of WT, CgA-KO and CST-KO mice. Data were analyzed by one-way ANOVA followed by Sidak’s multiple comparison test. n=8 mice per group (Statistical Power = 85%). *p<0.05, **p<0.01, ***p<0.001, ****p<0.0001

## 3 Conclusion and discussion

Our comprehensive longitudinal study underscores the pivotal role of CgA and CST in maintaining pancreatic homeostasis, including both structural and functional components. CgA is a unique pro-hormone containing counter-regulatory motifs. Among these, pancreastatin (PST; hCgA_250−301_) [48] has been reported to exert pro-diabetic, pro-inflammatory, and obesogenic effects [17], whereas CST (hCgA_352−372_) [49] exhibits anti-diabetic [19,37], anti-inflammatory [19,37], and anti-obesogenic [18] properties. Consequently, the simultaneous absence of both PST and CST in CgA-KO mice results in heightened insulin sensitivity, as previously documented in both normal chow diet (NCD) [38] and diet-induced obese (DIO) mice [17]. Supplementation of CgA-KO mice with PST led to increased glucose excursion during GTT and elevated glucose production during PTT, confirming PST as a pro-gluconeogenic, insulin-resistant peptide. Conversely, CST supplementation reduced glucose excursion in GTT and inhibited glucose production during PTT, highlighting CST as an insulin-sensitizing peptide as per our previous report in DIO mice [19]. Co-supplementation of CgA-KO mice with equimolar concentrations of PST and CST revealed that CST overrides PST’s gluconeogenic effect, suggesting CST’s dominance in regulating hepatic glucose output. Moreover, we have shown that CgA-KO mice maintain enhanced insulin sensitivity even after four months on a high-fat diet (60% calories from fat) [17]. CST’s beneficial effects on insulin sensitivity in DIO mice are mediated via suppression of monocyte-derived macrophage infiltration [19], attenuation of gluconeogenesis [19], and reduction of endoplasmic reticulum stress [37].

Morphometric studies demonstrating decreased beta cell numbers in CgA-KO and CST-KO mice align with biochemical findings of reduced insulin levels in these models. In contrast, increased alpha cell numbers in CgA-KO and CST-KO mice do not fully correlate with glucagon levels - no change in CgA-KO mice and decreased glucagon in CST-KO mice compared to WT mice. Intriguingly, despite elevated catecholamines, increased ketone bodies, and muscle insulin resistance, CgA-KO mice maintain euglycemia due to enhanced hepatic insulin sensitivity, as evidenced by GTT, insulin tolerance test, PTT and hyperinsulinemic euglycemic clamp studies [38]. The absence of the pro-gluconeogenic PST in CgA-KO mice likely contributes to this increased hepatic insulin sensitivity. Conversely, CST-KO mice allowed PST to exert its gluconeogenic effect without intervention by CST made CST-KO mice, which still express PST, develop insulin resistance due to unopposed PST action. Our current data reinforce this concept, as CST supplementation effectively dominates PST in regulating glucagon dynamics and gluconeogenesis.

We also observed discrepancies between *in vivo* GTT (15 min exposure) in intact animals and *ex vivo* GSIS (60 min exposure). Notably, glucose injections failed to stimulate *in vivo* insulin secretion in CST-KO mice but induced ∼29-fold insulin release *ex vivo*. This *in vivo* failure to respond to glucose with insulin secretion is likely attributable to elevated plasma catecholamines: norepinephrine (NE: ∼1.6-fold increase ) and epinephrine (EPI: ∼1.8-fold increase) in CST-KO mice [21]. Catecholamines are known to inhibit insulin secretion by activating α₂-adrenergic receptors on beta cells [50,51]. The α_2A_-adrenoceptor, the predominant subtype in mouse and human islets, reduces beta-cell excitability and insulin release by lowering cAMP levels and desensitizing of late exocytotic step(s) to cytosolic Ca^2+^ [50], leading to glucose intolerance *in vivo*. However, isolated islets studied *ex vivo* are free from catecholaminergic inhibition, enabling robust insulin secretion at high glucose. Thus, CST deficiency results in elevated catecholamines that potently inhibit insulin secretion, a constraint removed under *ex vivo* conditions.

GABA is linked to T1D through the autoantigen GAD65 and is thought to increase beta cell content in the islet [52]. Lower GABA levels might be one reason for the detected decrease in insulin producing cells in the CgA-KO mouse islets. However, the differences seen in CST-KO islets are probably caused by other mechanisms affecting insulin content or beta cell numbers since the GABA and histamine values upon CST KO are similar to WT islets. Beta cells contain highest concentrations of polyamines (putrescine, spermidine, and spermine) [53,54], where they regulate proinsulin biosynthesis and secretion of insulin [55]. The depletion of polyamines in isolated mouse islets has been associated with impaired glucose-stimulated insulin secretion, insulin content, insulin transcription, and DNA replication [56,57]. Polyamine levels were reported to be diminished in aging and obese mice [58], which are resistant to insulin. CST-KO mice have also been reported to be insulin resistant [19]. The markedly decreased spermine levels in CST-KO pancreas might be the reason for the observed increased insulin plasma levels and/or resistance to insulin in CST-KO mice. Also the observed lower spermine levels in the CgA-KO and CST-KO mice can result in disturbed insulin secretion [44] and less uptake of Ca^2+^ by the beta cells [59]. In obese mice the spermine to spermidine ratio is even similarly disturbed in the pancreatic islets as in our knockout mice [58].

Altogether, more research is necessary to identify the pathway(s) resulting in the affected islet composition in the CgA-KO and CST-KO mice. With our unique methodology, we were able to spatially investigate the pancreatic microenvironment - a crucial step towards identifying and understanding the pathways affecting islet composition in knockout mice. However, since our current methodology is insufficient to detect enzyme expression and activity, future research should focus on studying the enzymes regulating catecholamine degradation and spermine pathways to clarify the underlying mechanisms of the pancreatic microenvironment further.

The CgA cleavage product CST is increasingly implicated in metabolic regulation, including insulin sensitivity, glucose homeostasis, and inflammation. The enhanced beta cell responsiveness in isolated islets of CST-deficient mice suggests a physiological role for CST in restraining insulin hypersecretion, possibly to prevent beta cell exhaustion in metabolic stress conditions. The discrepancy between *in vivo* GTT and *in vitro* GSIS, where isolated islets from CST-KO mice were highly responsive to glucose *in vitro*, but not *in vivo*, may be partially explained by the hyperadrenergic state of these mice. CST normally acts as a physiological brake on catecholamine release, and the increased NE and EPI levels may thus limit insulin secretion from islet beta cells. Thus, modulating CST activity could offer a novel therapeutic approach for metabolic disorders characterized by insulin resistance, such as T2D and obesity. Specifically, CST antagonists may enhance endogenous insulin output in the context of beta cell insufficiency, while CST mimetics/analogs could be beneficial where insulin hypersecretion contributes to disease progression.

Results of this study should also be interpreted with caution since the use of full body (systemic) KO models for CgA and CST has some limitations. Whole-body KO can introduce confounding effects from other tissues (e.g., liver, fat, muscle) or can be the result of developmental issues, which may also influence the observed metabolic phenotype. The development of a conditional CgA and CST knockout model that can be turned on and off in target tissues such as the pancreas would be very useful for future research. Human and mouse islets are slightly different in morphology and endocrine composition [60], therefore the spatial MS, GSIS and fluorescent staining of this study should be repeated on CgA or CST stimulated human spheroids or islets to confirm the effects in human. Additionally, future research should focus on further profiling of macrophages and other immune cell populations to clarify their role in neuro-immunomodulation in the context of CgA and CST absence. Surprisingly, we didńt observe significant differences of catecholamine levels in the pancreas, while catecholamine plasma levels of the KO mice are increased. However, we didńt look into NE or EPI degradation products, future research should confirm if the catecholamine degradation is indeed affected upon CgA or CST KO.

Recent studies have highlighted the crucial connection between the local immune system in the pancreas and the peripheral nervous system in the development of autoimmune diseases [61–63]. Interfering with pancreatic nerve signals through surgery, chemical blockage, or electric stimulation has been shown to prevent the onset of T1D in mice [61–63] Additionally, depleting macrophages during the onset of autoimmune diabetes in mice can halt T1D onset [62]. These findings indicate that the connection between the local immune system in the pancreas and the peripheral nervous system plays an important role in the development of autoimmune disease. This connection may involve NE signaling, where NE is produced by neurons but locally regulated by nerve-associated macrophages, potentially through β_2_ adrenergic signaling, as observed in hypertension [64]. This mechanism may explain how CgA and CST influence autoimmune disease development since 1) NE/EPI levels are high in CgA-KO and CST-KO mice, 2) both KO mice are characterized with hypertension and, 3) CgA and CST levels are elevated in autoimmune diseases. Clarification of this link between CgA/CST and autoimmune disease development through catecholamine signaling among nerves, immune cells, and endocrine cells could provide novel insights into therapeutic strategies for autoimmune diseases in the future.

Altogether, our findings delineate the opposing roles of CgA cleavage products PST and CST in orchestrating pancreatic and peripheral glucose homeostasis, highlighting a neuro-immune-endocrine regulatory axis (Fig 9). Our PTT data establish CST’s dominance over PST in suppressing gluconeogenesis, underscoring the therapeutic potential of modulating CgA-derived peptides. By targeting the neuro-immune-endocrine (structural and functional components) axis, through CST-based therapy or PST inhibition could represent a novel strategy to improve insulin sensitivity and combat diabetes and related metabolic disorders. This study provides a foundation for precision peptide therapeutics aimed at complex metabolic and autoimmune diseases.

**Fig. 9.**
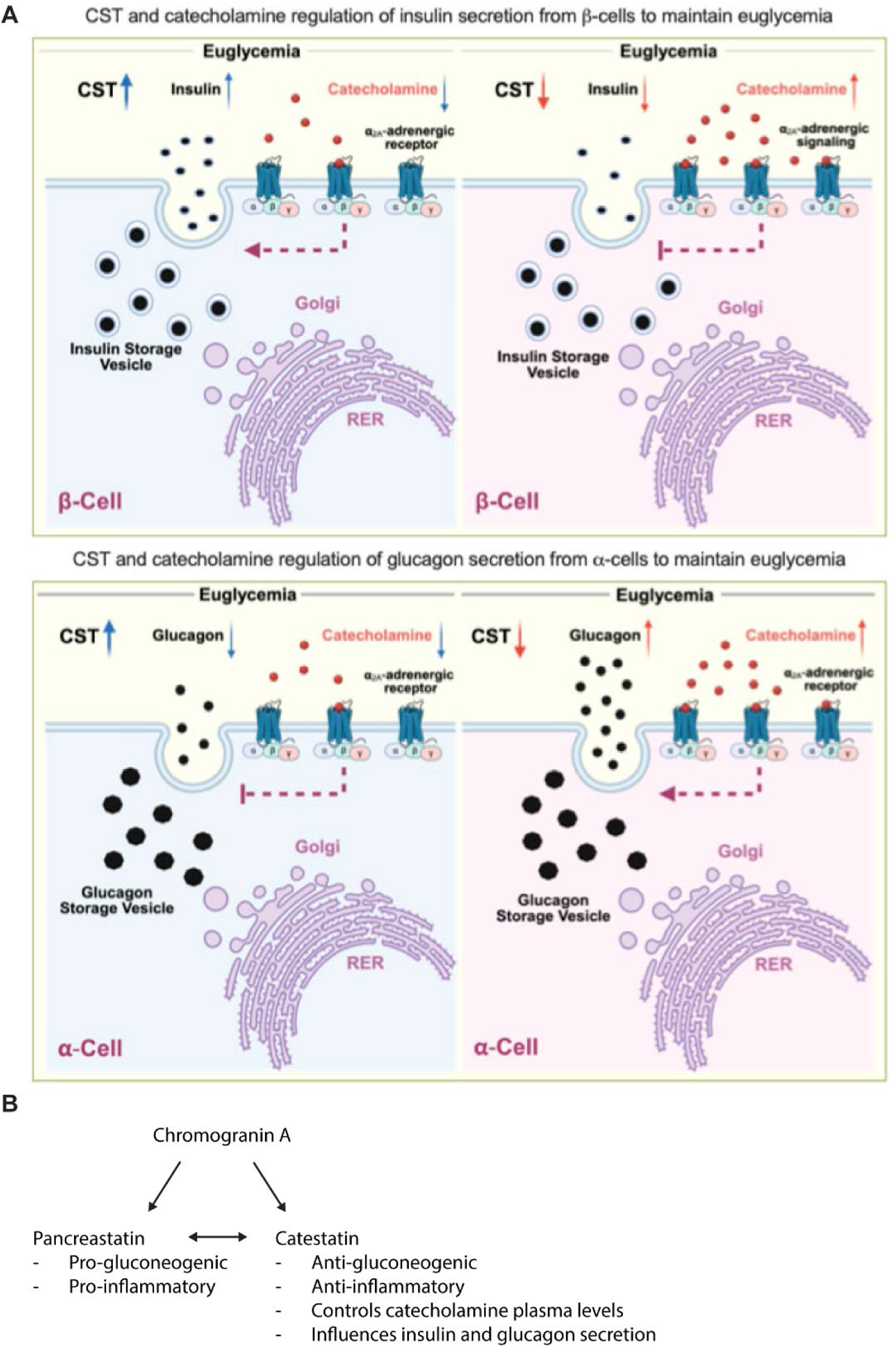
Proposed working mechanism of CgA and CST on pancreatic homeostasis. **A)** Proposed working mechanism of euglycemia in the presence and absence of catestatin. **B)** Summarized effects of chromogranin A cleavage products catestatin and pancreastatin.

## 4 Material and Methods

### 4.1 Human blood samples for CST ELISA

The collection of blood samples was approved by the Regional Research Ethical Committee in Uppsala (Dnr 2014/485) and was conducted in consistency with The Declaration of Helsinki. All participants gave their written informed consent prior to inclusion in the study. CST levels in plasma were determined using ELISA (CUSABIO CSB-E17355h). Patient characteristics are listed in S-Table 1.

### 4.2 Mice

Male wild type (WT), CgA-knockout (KO) mice and CST-KO mice (5-8 months old) on C57BL/6 background were kept in a 12 hours dark/light cycle on normal chow diet (NCD: 13.5% calorie from fat; LabDiet 5001, Lab Supply, Fort Worth, TX). These mouse studies were approved by the UCSD and Veteran Affairs San Diego Institutional Animal Care and Use Committees and conform to relevant National Institutes of Health guidelines. Organs were harvested after deeply anesthetizing the mice with isoflurane followed by cervical dislocation. For the immunoblotting and immunofluorescent experiments, organs (adrenal gland, pancreas, bone marrow) were harvested from male C57BL/6J mice (Taconic, Denmark) or non-obese diabetic (NOD) mice. These studies were approved by the Regional Animal Ethics committee in Uppsala, Sweden (permit no: 5.8.18-01462/2023, 5.8.18-01540/2024). All animal experiments were conducted according to local standard operating procedures regarding humane endpoints.

### 4.3 Tissue preparation for immunohistochemistry

Pancreata were harvested from WT, CST -KO, and CgA-KO mice. Pancreata were cut in two parts and snap frozen. For immunostainings, the pancreata were embedded in O.C.T, sectioned in 10 µm sections at -20℃ and captured on microscope glass slides (Epredia; J1830AMNZ). For mass spectrometry imaging, the frozen pancreas tissues were sectioned at a thickness of 12 μm using a Leica CM3050S cryostat set at -20°C, thaw-mounted on stainless steel MALDI-target plates and subsequently stored at -80°C until further analysis.

### 4.4 Fluorescent staining of the pancreatic islets

Pancreatic sections were fixed for 10 min in 4% paraformaldehyde (PFA) followed by washes in phosphate-buffered saline with 0.1% Tween (PBST) (Medicago; 274713; SIGMA-ALDRICH; SZBA3190V). Afterwards slides were treated for 10 min with 0.2% Triton X100, followed by 30 min in blocking solution (PBS supplemented with FBS 2%, saponin 0.2%, NaAz 0.1%, 1:1000 FC blocker (BD Bioscience; 553142)). To prevent antibody cross-reactivity, the insulin staining was performed first. To do so, slides were incubated o.n. (overnight) at 4°C with guinea pig insulin primary antibody in blocking buffer (1:1000, Fitzgerald; 20-IP35). The next day, slides were washed three times with PBST for 5 min followed by staining for 30 min at r.t. with the secondary antibody anti-guinea pig-488 (1:500, Invitrogen; A-11073). After washing three times, slides were incubated for 1 hour at r.t. with either mouse anti-glucagon 1:200 (1:200, Proteintech; 67286-1-Ig) and rabbit anti-somatostatin (1:500, Abcam; ab111912) or rabbit anti-norephrine transporter (1:200, Abcam; ab254361) and rat anti-IBA1 (1:1000, Synaptic Systems; 234017) or rabbit anti-Tyrosine Hydroxylase (Abcam; ab137869) or 488-anti-mouse CD3 (1:300, Biolegend; 100210) and rabbit anti-CST (1:1000, Proteintech; 289-MM-0288). Slides were washed three times with PBST for 5 min followed by staining for 30 min at r.t. with anti-mouse 555 (1:1000, Invitrogen; A21127) and anti-rabbit-647 (1:2000, Invitrogen; A31573) or anti-rabbit 555 (1:1000, Invitrogen; A31572) and anti-rat-647 (1:1000, Invitrogen; A48272). After washing three times with PBST, the slides were stained for 10 min with Hoechst (1:10.000, ThermoFisher; 33342) to visualize the nuclei. Afterwards, slides were washed three times with PBS. After the final wash the cover glass (24×50 mm, VWR; 631-0147) was applied using ProLong™ Gold Antifade Mountant (ThermoFisher; P36934). For imaging, the slide scanner Zeiss Axio Scan Z1 with a 20x objective was used.

### 4.5 Analysis of insulin, glucagon and somatostatin cells in the pancreatic islet

Slide scanner pictures were analysed using QuPath [65] (S-Method Fig. 1). Islets were annotated based on staining in all channels visualizing the complete islet. The machine-learning software was trained on WT pancreas images to detect cell number per islet using the Hoechst nuclei staining channel and the following nucleus parameters: background radius: 15 px; median filter radius: 0 px; sigma: 3 px; minimum area: 10 px^2; maximum area: 1,000 px^2; intensity parameters: threshold 100; cell parameters: cell expansion 5 px. The obtained cell classifier was used to determine islet shape features (area, length, circularity, solidity, maximum diameter, minimum diameter, and nucleus/cell area ratio) and to calculate the islet density (islet density= islet area/pancreatic section area). This was followed by obtaining the numbers of insulin, somatostatin, and glucagon-producing cells per islet. Afterwards, insulin, somatostatin and glucagon cell ratios were calculated: Positive cells % per islet = (100/total cell number) *target cell number).

### 4.6 *In vivo* glucose tolerance test (GTT)

Mice were fasted overnight for eight hours before starting the GTT. Blood glucose was measured from tail tip blood drops using a glucose strip (True Metrix, total diabetes, Whitestown, IN) and a glucometer (total diabetes, Whitestown, IN) after 15, 30, 60, 90 and 120 min of administration of intraperitoneal dextrose (2 g/kg). Data is presented as blood glucose concentration (mg/dL) over time and as are under the curve above baseline.

### 4.7 Pancreatic islet isolation and islet glucose-stimulated insulin secretion (GSIS)

Pancreatic islets were isolated from mice by injecting collagenase P solution (1.4 mg/mL; #11213873001, Roche) into the pancreas via the common bile duct. The pancreas was subsequently incubated at 37 °C with intermittent vortexing (10 seconds at medium speed) until fully digested. Enzymatic activity was quenched by adding ice-cold RPMI medium supplemented with 10% fetal bovine serum (FBS), followed by gentle agitation. The digested tissue was centrifuged at 300 × g for 3 minutes at 4 °C. The supernatant was discarded, and the pellet was washed twice with cold complete medium, then passed through a 500 µm filter (#U-CMN-500-B, Component Supply). The filtered suspension was centrifuged again and resuspended in a smaller volume to facilitate loading onto a 70 µm cell strainer (#352350, Corning). The strainer was then inverted into a 60 × 15 mm tissue culture dish (#83.3901.500, SARSTEDT) containing 5 mL of complete RPMI medium. Islets retained on the membrane were collected under a stereomicroscope and cultured in RPMI containing 11.1 mM glucose, 10% FBS, 100 U/mL penicillin– streptomycin, and 2 mM GlutaMAX. Following isolation, the following parameters were assessed: the number of islets per mouse, insulin content per 10 islets, glucose-stimulated insulin secretion (GSIS), GSIS normalized to insulin content and the insulin secretion index.

### 4.8 *In vivo* insulin and glucagon measurements

All mice were fasted for eight hours before starting the glucose injections. Blood (50 µl) from WT, CgA-KO and CST-KO mice was collected at 0 min from the tail tip using a capillary tube and kept on ice. Mice were injected with dextrose (2g/kg intraperitoneally) and blood (50 µl) was collected again after 15 min of dextrose administration. Blood samples were centrifuged at 12,500 RPM for 10 min at 4°C to separate the plasma. Plasma was stored at -80°C until analysis of insulin and glucagon levels. 25 µl plasma was used for insulin and glucagon assay using MSD U-PLEX ELISA kit (Meso Scale Diagnostics, Rockville, MD).

### 4.9 Pyruvate Tolerance Test (PTT)

All mice were fasted for eight hours before starting the PTT. Blood glucose was measured on a blood drop collected from the tail tip using a glucose strip and a glucometer at 0 min. Sodium pyruvate (2 g/kg intraperitoneally) was injected, followed by blood glucose measurements from tail tip using a glucose strip and a glucometer after 15, 30, 60, 90 and 120 min after administration. Data is presented as blood glucose concentration (mg/dL) over time.

### 4.10 Mouse cytokine and hormone measurements

25 μl mouse plasma was used in a customized U-PLEX Custom Metabolic Group Assay kit (Meso Scale Diagnostics, Rockville, MD) by following the manufacturer’s protocol. Mouse hormones and cytokines were measured: glucagon, insulin, KC/GRO, IL-1β, IL-6, IL-10, MCP-1 (CCL2) and TNF-α, using a MESO SECTOR S 600MM Ultra-Sensitive Plate Imager. Insulin and glucagon levels were presented as ng/mL. Cytokines and chemokines were presented as pg/mL.

### 4.11 Measurement of Plasma Catecholamines

Plasma catecholamines were quantified using an Atlantis dC18 column (100 Å, 3 µm, 3 × 100 mm; Waters Corp., Milford, MA) on an ACQUITY UPLC H-Class System equipped with an electrochemical detector (ECD model 2465; Waters Corp.) as previously described [21,66]. The isocratic mobile phase consisted of phosphate–citrate buffer and acetonitrile (95:5, v/v) at a flow rate of 0.3 mL/min. For extraction, 2 ng of 3,4-dihydroxybenzylamine (DHBA) was added as an internal standard to 100 µL of plasma. Catecholamines were adsorbed onto ∼15 mg of activated aluminum oxide by rotating the mixture for 10 minutes. Following adsorption, the aluminum oxide was washed with 1 mL of ultrapure water, and catecholamines were eluted with 100 µL of 0.1 N HCl. The electrochemical detector was set at a sensitivity of 500 pA for catecholamine detection. Chromatograms were analyzed using Empower software (Waters Corp.). Catecholamine concentrations were normalized to the recovery of the internal standard and expressed as ng/mL.

### 4.12 Immunoblotting

Macrophages grown from mouse bone marrow [67], or tissue (medulla, isolated pancreatic islets [68]) were lysed in lysis buffer (1% SDS, 10 mM TrisHCl, pH 6.8). Protein concentration was determined according to manufacturer’s instructions (Bio-Rad; 500-0114). Afterwards, for each condition 60µg of protein was loaded and run on 10 % mini-protein TGX gels (Bio-Rad; 4561033) in Tris/Tricine/SDS buffer (Bio-Rad; 1610744) followed by transfer for 60 min at 100V at 4°C in Tris/Glycine buffer (Bio-Rad; 1610771) with methanol (Supelco; 1263283323) using a PVDF membrane. The membranes were taken out the cassette and washed with distilled water, followed by blocking in 3% BSA buffer for 1 h at r.t.. Membranes were stained o/n at 4°C with primary antibody rabbit anti-chromogranin A (1:1000, Invitrogen; PA5-35071) or rabbit anti-catestatin (1:1000, Proteintech; 289-MM-0288). Next day, the membranes were washed three times quickly and three times for 10 min with TBS-t 0.02%. Followed by staining with secondary goat-a-rabbit antibody 1:5000 dilution (IRDye800; P/N 925-32211) for one hour at r.t. Afterwards, the membrane was again washed 3 times with TBS-t 0.02%. Both membranes were scanned on a Bio-Rad ChemiDoc MP imager to visualize the fluorescent staining.

### 4.13 Spatial Mass Spectrometry

Two consecutive 12 µm pancreatic sections were taken containing tissue of 3 WT, 3 CgA-KO and 3 CST-KO mice. One section was stained with hematoxylin-eosin (HE) (S-Method Fig. 2) to annotate the pancreatic islets using QuPath. The other slide was used for neurotransmitter analysis of serotonin (5-HT), gamma-aminobutyric acid (GABA), histamine, cysteine, taurine, creatine, spermidine, histidine, NE, tyrosine, spermine and hypoxanthine. Derivatization matrix FMP-10 (4.4 mM in 70% acetonitrile, Tag-ON AB, Uppsala, Sweden) [69] (S-Table 2) was applied with TM-Sprayer (HTX-Technologies, Chapel Hill, NC, USA). The spraying method was set up to include 30 passes, with a flow rate of 80 μl/min at a temperature of 80°C. The nozzle velocity was adjusted to 1100 mm/min, while the track spacing was set at 2.0 mm, and N2 was set 6 psi. Full tissue MSI experiments were performed using a timsTOF fleX MS imaging instrument in positive ion mode (Bruker Daltonics GmbH, Bremen, Germany). Online calibration was performed using m/z 555.2231, an abundant ion cluster signal of FMP-10. Data acquisition was setup in single scan mode at 30 µm lateral resolution, collecting 200 shots per pixel in the m/z range 300–1000 using flexImaging 5.0 and timsControl 6.0 (Bruker Daltonics). Islets were mapped on the tissue with 5-HT signal and overlay with stained HE staining. For each islet a corresponding ROI of exocrine tissue was drawn. Metabolite identities were validated by MS/MS directly from tissue and compared with MS/MS of FMP-10 derivatized analytical standards (S-Method Fig. 3&4). MS/MS data was collected on a MALDI-FTICR MS (solariX, Bruker Daltonics). The isolation window for fragmentation was set to 1 Da and collision energy of 35 eV.

### 4.14 Analysing neurotransmitters in the islets and exocrine pancreas

Islet selections in Qupath were exported using the SCiLS lab extension (Bruker Daltonics) and imported into the flexImaging (Bruker Daltonics, Bremen, v.5.0) software. To match and optimize the pancreatic islet location from the HE staining, the serotonin signal was used. After identifying the pancreatic islets, annotations were made for the matching exocrine tissue (serotonin negative area). Expression data of neurotransmitters in the pancreatic islet and exocrine tissue was extracted and visualized in graphs. Illustration showing islet selection and data extraction can be found in the supplementary material (S-Method Fig. 2).

### 4.15 Statistics and Reproducibility

Data are expressed as mean ± SEM, all datapoints are shown by dots and ‘n’ in the figure legends represents the biological replicates of the group. One-way or two-way ANOVA with Sidak’s post-hoc tests or non-parametric Mann-Whitney test were applied for multiple comparisons. A p-value of less than 0.05 was considered statistically significant. Sample sizes, statistical analyses, and statistical power are indicated in the corresponding figure descriptions.

## 6 Data availability statement

The datasets generated and analysed in the current study are available in the FigShare repository (https://figshare.com/s/8dace17885070d6916f1, DOI: 10.17044/scilifelab.28172552). Raw spatial MS data, slide scans and human data is available upon request via Gustaf Christoffersson.

## 7 Author Contributions

S.K.M. was responsible for mouse husbandry, peptide treatments, and harvesting the pancreata. He conducted GTT, *in vivo* hormone measurements, and PTT experiments with help from K.T., analyzed the data and made the graphics. K.T. and M.R. ran the MSD kit on the mouse samples. S.K. performed H&E staining on histological slides. S.K. and S.J. ran Western blot on pancreatic islets. E.M.M., H.I., D.E. and M.B. have performed H&E stainings, fluorescent stainings and image analysis. T.V.R. performed and analysed the islet GSIS. D.E. performed immunoblotting. M.N. and A.N. sectioned, prepared and ran the pancreatic slices in the Maldi-MS. M.N., A.M.N., A.A.M. and E.T.J. performed data analysis. D.E. provided human plasma samples and K.S. provided NOD mice material. S.J. made the working hypothesis cartoon. E.M.M. assisted with data interpretation and compiled the manuscript. G.C., P.E.A., S.M.K and E.T.J. provided expertise. The manuscript was written by E.M.M., G.C., and S.K.M. All authors contributed in writing and editing the manuscript.

