## Supplementary Material for "Chromogranin A and catestatin regulate pancreatic islet homeostasis, endocrine function, and neurotransmitter signaling"

Supplementary Tables: 2
Supplementary Figures: 14
Supplementary Method Figures: 4

**S-Table 1. Characteristics of healthy and T1D participants.**

| Diabetes onset (year) | Gender (M/F) | Age (year) | BMI ($kg/m^2$) | C-peptide (nmol/l) |
|---|---|---|---|---|
| 1978 | M | 42 | 27 | <0,01 |
| 1999 | M | 39 | 28,7 | <0,01 |
| 1989 | M | 43 | 29,3 | <0,01 * |
| 1997 | M | 35 | 25,1 | <0,01 |
| 1991 | M | 30 | 27,2 | <0,01 |
| 2005 | F | 21 | 22,9 | <0,01 |
| 1999 | M | 45 | 27,7 | |
| 1990 | F | 43 | 22,3 | |
| 1988 | F | 42 | 26,8 | <0,01 |
| 1995 | M | 27 | 30,7 | <0,01 |
| 2010 | M | 24 | 22,9 | <0,01 |
| 1995 | M | 38 | 24,8 | 0,05 * |
| 2001 | M | 49 | 26,3 | <0,01 |
| 1997 | M | 35 | 25,6 | 0,04 |
| 2003 | F | 23 | 24,2 | <0,01 |
| 1987 | M | 35 | 24,2 | <0,01 |
| 1994 | F | 37 | 25,1 | <0,01 |
| 1997 | M | 40 | 22,6 | <0,01 |
| 2001 | F | 25 | 25,4 | <0,01 |
| 2002 | F | 25 | 23,3 | <0,01 |
| 1999 | M | 45 | 27,3 | 0,4 |
| 1994 | F | 41 | 30,5 | 0,05 |
| 1998 | M | 33 | 35,5 | <0,01 |
| 1981 | M | 41 | 24,8 | 0,01 |
| Healthy | F | 51 | 27,1 | 0,5 |
| Healthy | F | 26 | 22,5 | 0,5 |
| Healthy | F | 26 | 20,7 | 0,5 |
| Healthy | F | 36 | 22,1 | 0,5 |
| Healthy | M | 39 | 22,8 | 0,4 |
| Healthy | F | 23 | 21,1 | 0,57 |
| Healthy | F | 25 | | |
| Healthy | F | 25 | | |
| Healthy | M | 26 | | |
| Healthy | M | 30 | | |
| Healthy | M | 23 | | |
| Healthy | M | 58 | 23,9 | 0,7 |
| Healthy | M | 26 | 21,7 | 0,4 |

Characteristics of individuals from whom blood samples were used in the CST ELISA in main Fig. 1. The table displays year of diabetes onset (or healthy), gender: male (M) or female (F), age in years, body mass index (BMI) expressed in units of kilograms (kg) divided by square of height in metres ($m^2$) and C-peptide levels in nanomoles (nmol) per litre (l).

**S-Table 2. Metabolites identified and validated in the pancreas.**

| Analyte name | Analyte short name | Theoretical m/z |
|---|---|---|
| Serotonin | 5HT TT | 444.2062 m/z ± 7.7 ppm |
| Gamma-Aminobutyric acid | GABA_H2O TT | 353.1642 m/z ± 8.1 ppm |
| Histamine | Histamine TT | 379.191 m/z ± 9.4 ppm |
| Cysteine | Cysteine TT | 389.1321 m/z ± 9 ppm |
| Taurine | Taurine TT | 393.1267 m/z ± 8 ppm |
| Creatine | Creatine TT | 399.1807 m/z ± 12 ppm |
| Spermidine | Spermidine TT | 413.2694 m/z ± 7.2 ppm |
| Histidine | Histidine TT | 423.1806 m/z ± 7.4 ppm |
| L-tyrosine | Tyrosine TT | 449.1851 m/z ± 10 ppm |
| Spermine | Spermine TT | 470.327 m/z ± 10 ppm |
| Hypoxantine | Hypoxantine TT | 404.1503 m/z ± 9 ppm |

The table lists analytes and their short name used in this study. Theoretical mass of the analyte is shown in (m) divide by charge number (z).

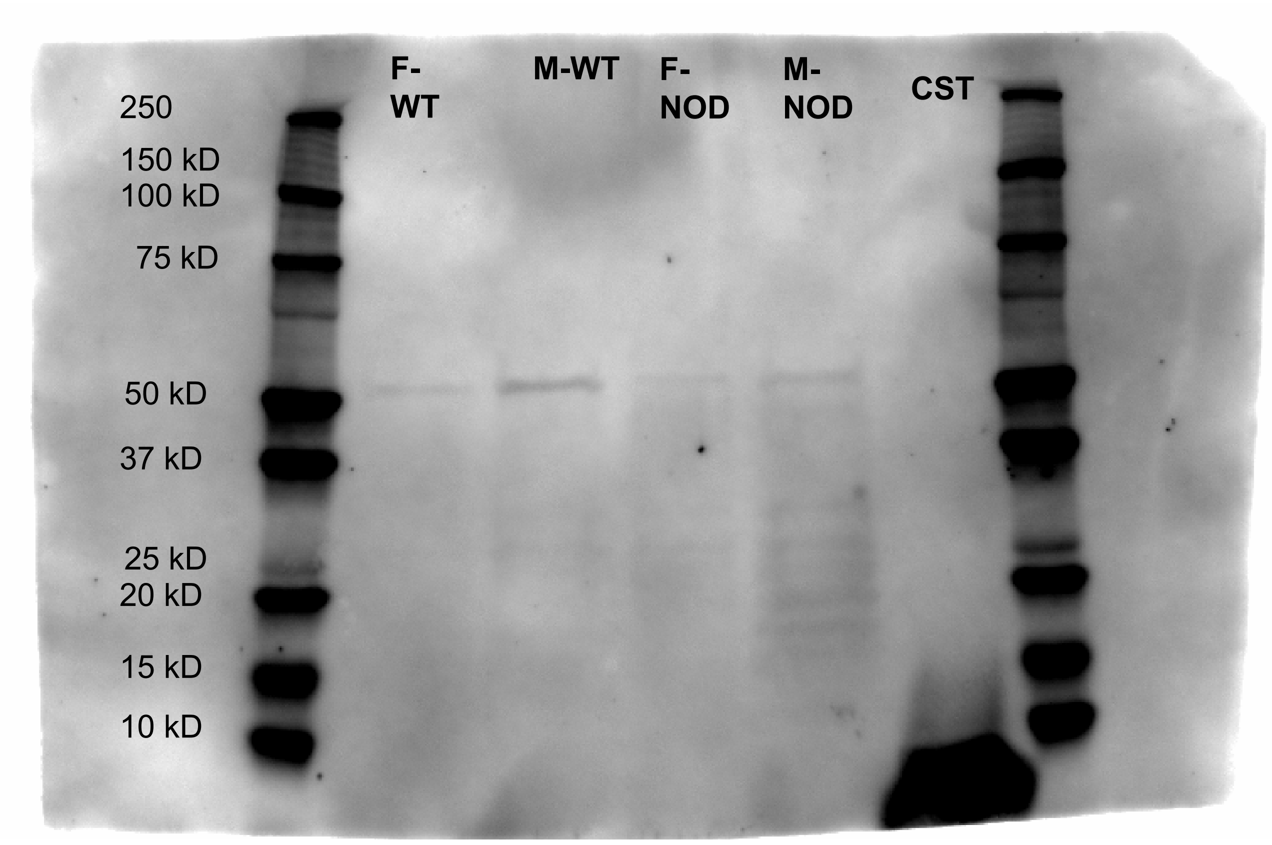


**CST staining**

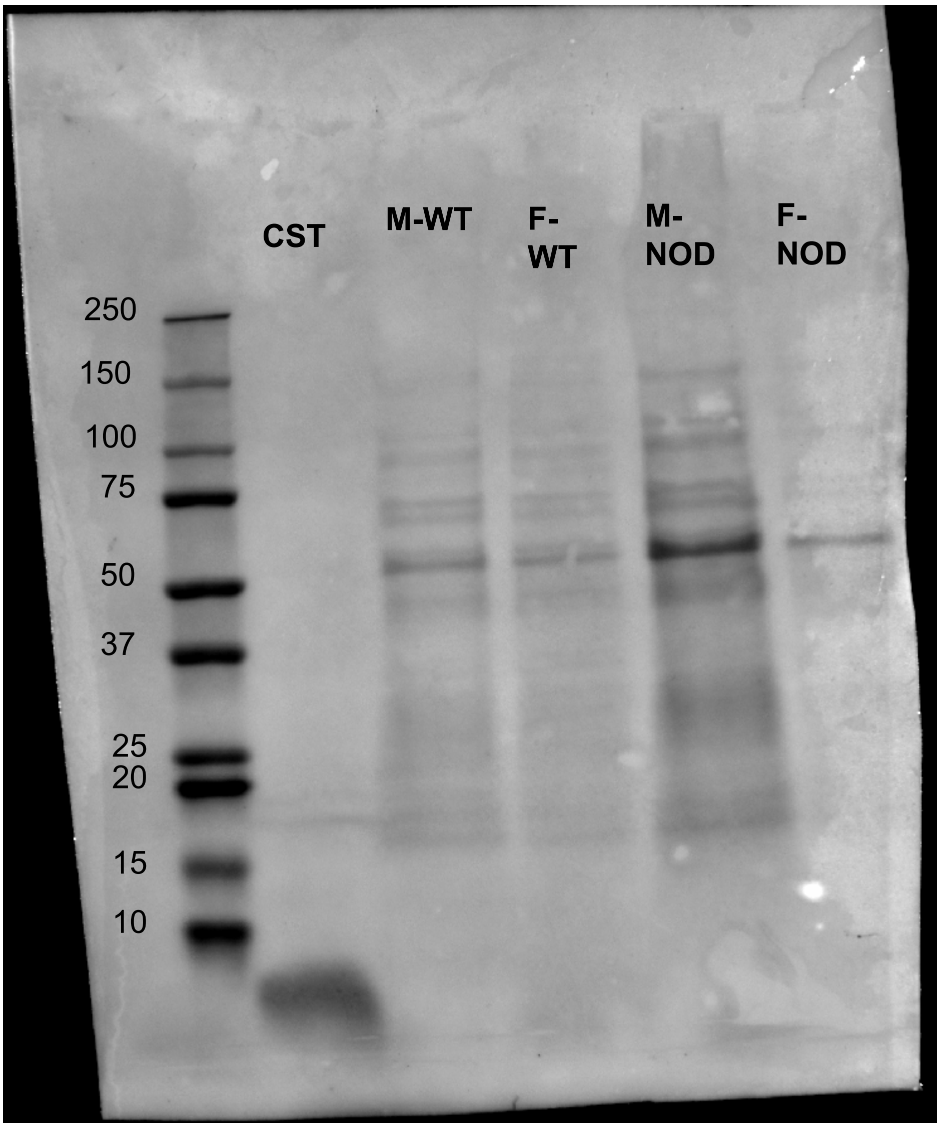


**Protein staining**

**S-Fig.1 Original immunoblot NOD mice from main Figure 1E.**
Original Western blot images showing catestatin staining or protein staining in wildtype (WT) or 8 week NOD mice (NOD) of Female (F) and Male (M) samples.
.

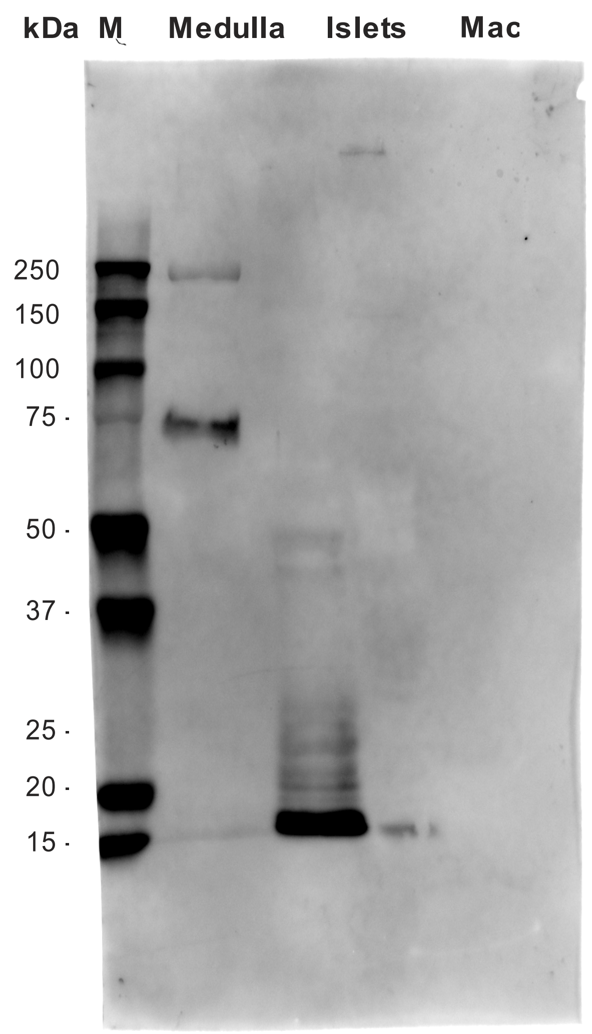


CgA staining

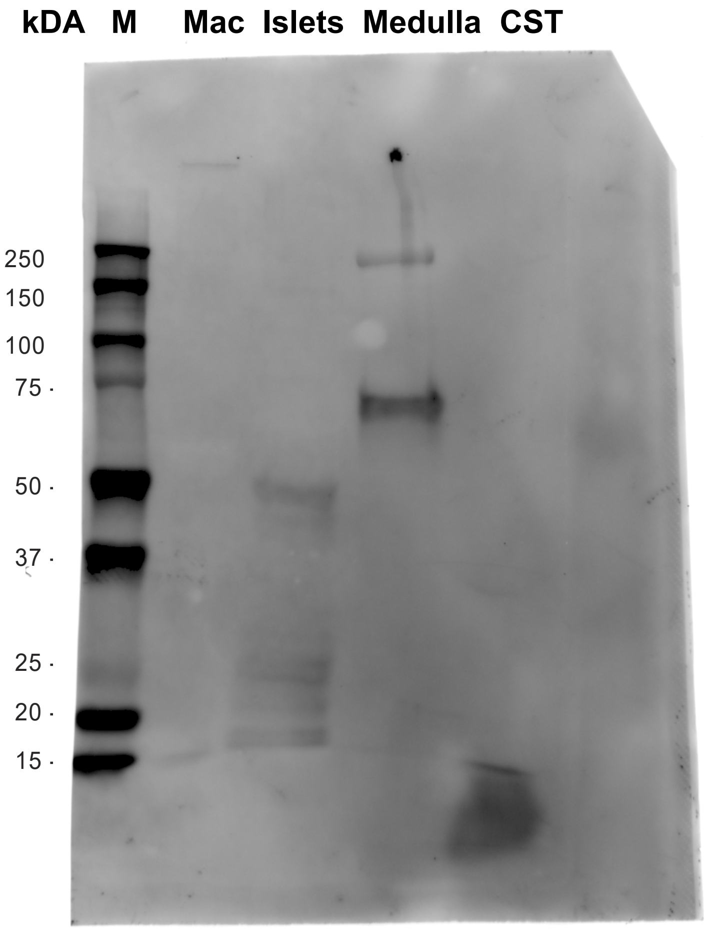


CST staining

**S-Fig.2 Original immunoblots from main Fig. 1F & G**
Original Western blot images showing chromogranin A or catestatin staining in the adrenal medulla, pancreatic islets and macrophages (mac).

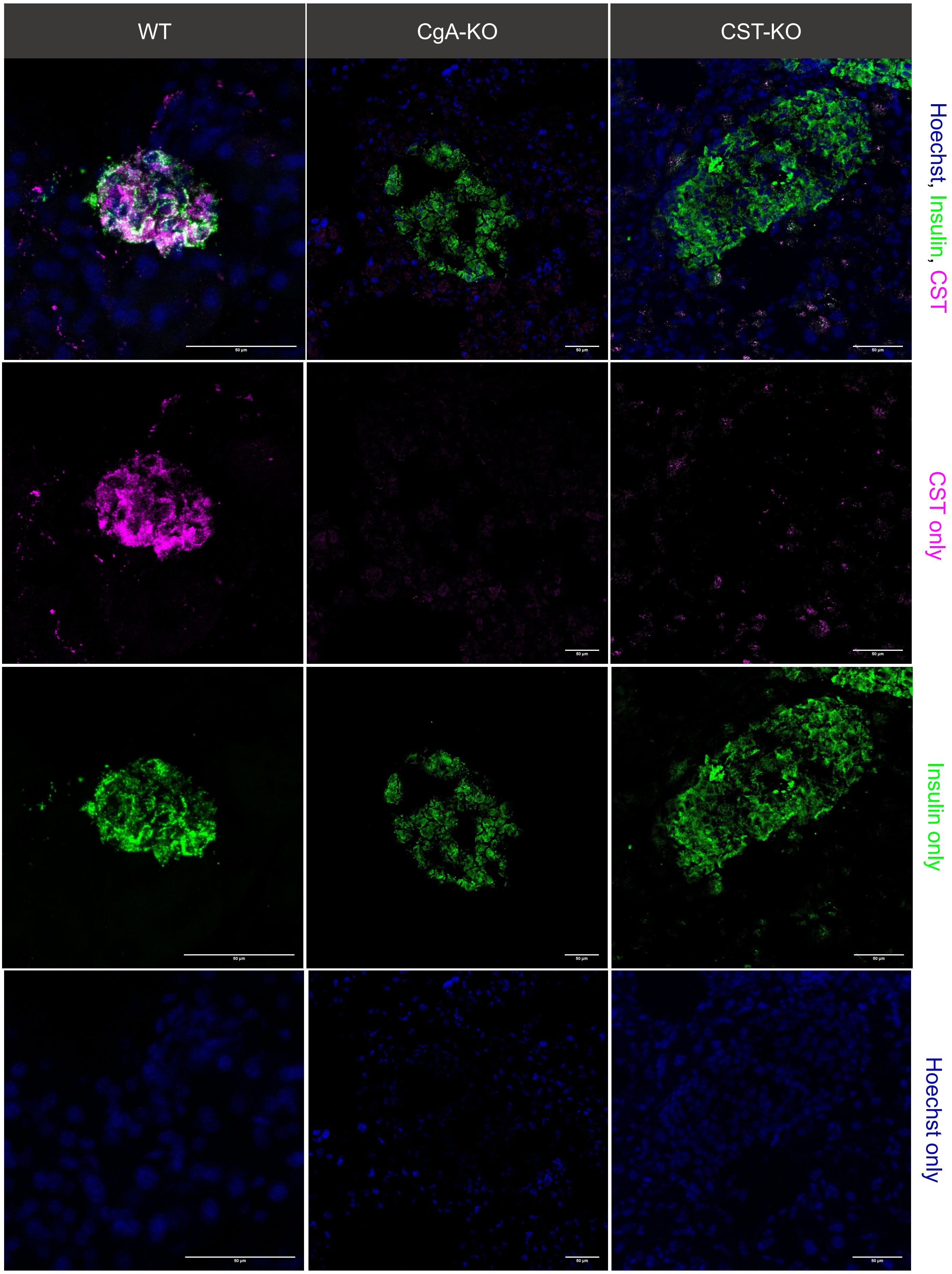


**S-Fig.3 Pancreatic islet CST staining confirming KO.**
Immunofluorescent staining for catestatin in WT, CgA-KO and CST-KO pancreas.

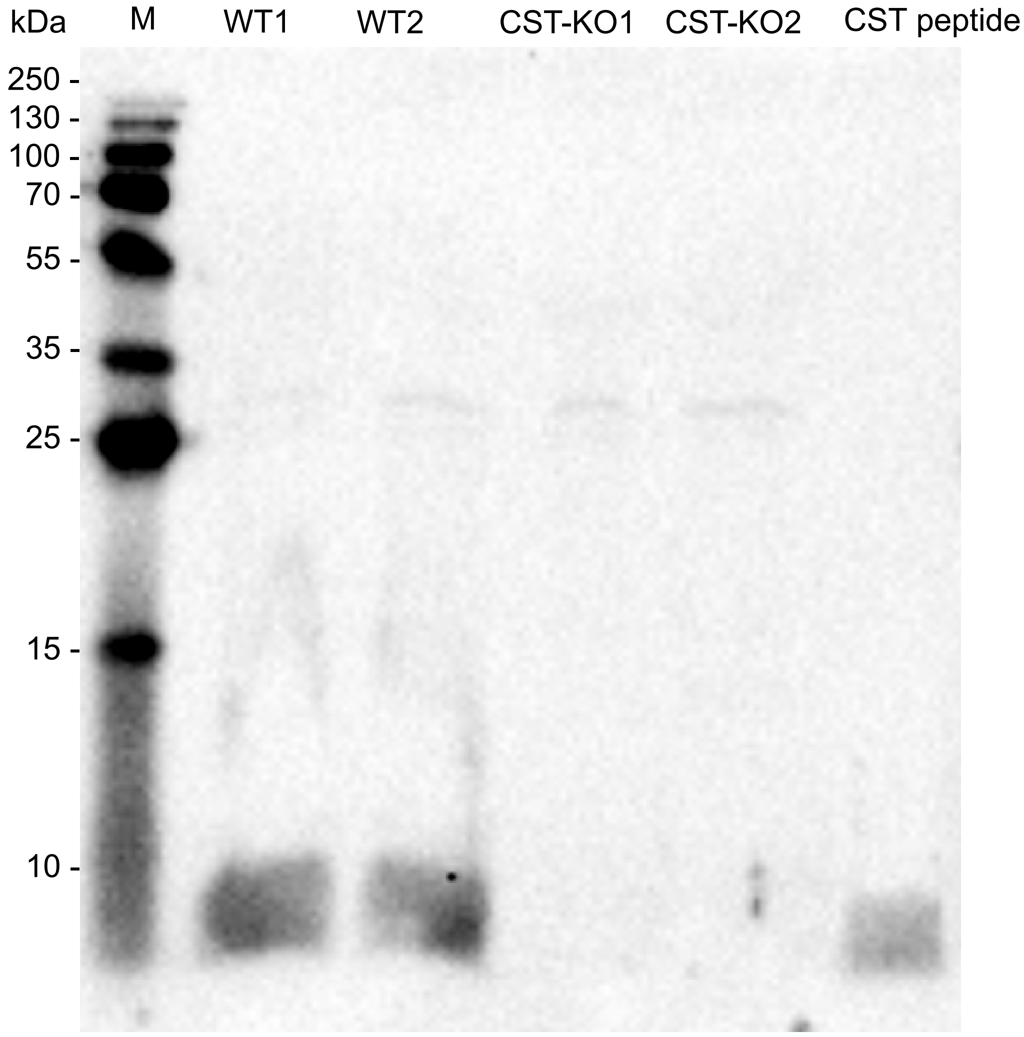


**Catestatin staining**

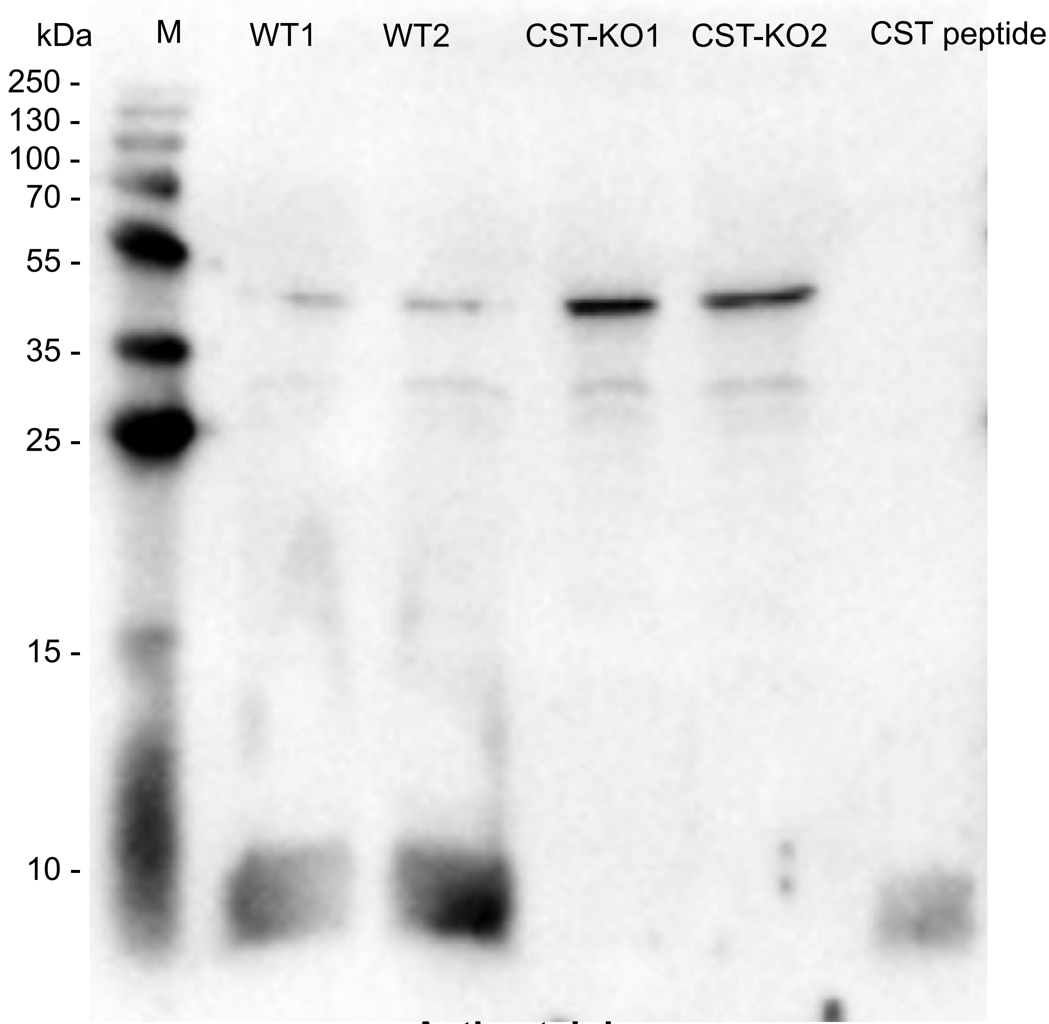


**Actin staining**

**S-Fig.4 Original immunoblots from main Fig. 2**

Original immunoblotting images showing catestatin (CST) or actin staining for WT mice, CST-KO mice or CST peptide only (positive control) including marker (M) and kilodalton (kDa).

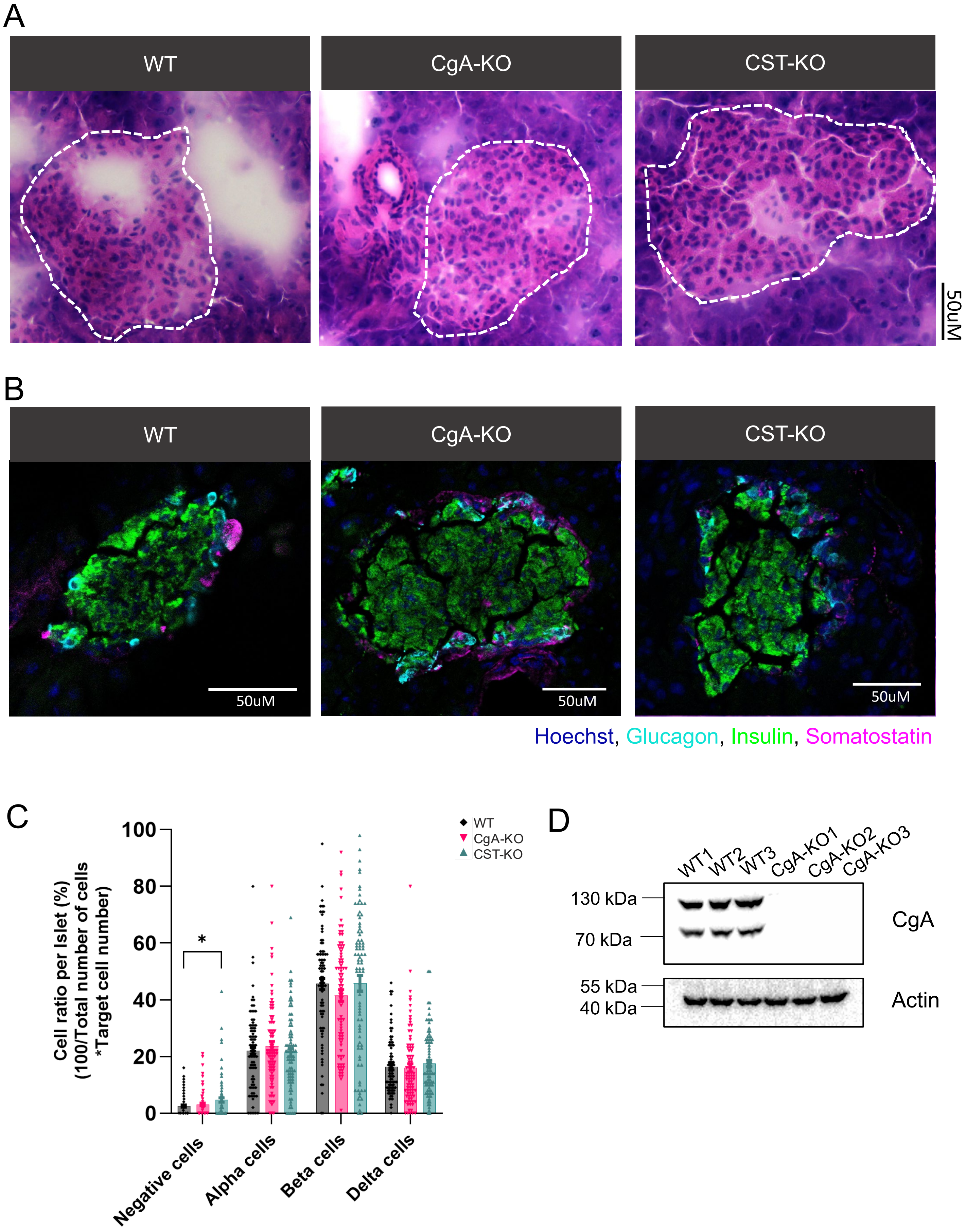


**S-Fig.5 Female pancreatic islet composition**

**A)** H&E staining of female WT, CgA-KO and CST-KO pancreatic slices including annotations for islets (white dotted lines). **B)** Representative images of immunofluorescent staining of glucagon (cyan), insulin (green), somatostatin (magenta) and Hoechst (blue) on WT, CgA-KO and CST-KO pancreatic slices. **C)** Quantification of alpha/beta/delta/negative cells per islet. Quantification is based on the staining displayed in panel B. N=3 mice per group. **D)** Immunoblotting images showing CgA and actin staining in WT, CgA-KO and CST-KO mice. *P<0.05

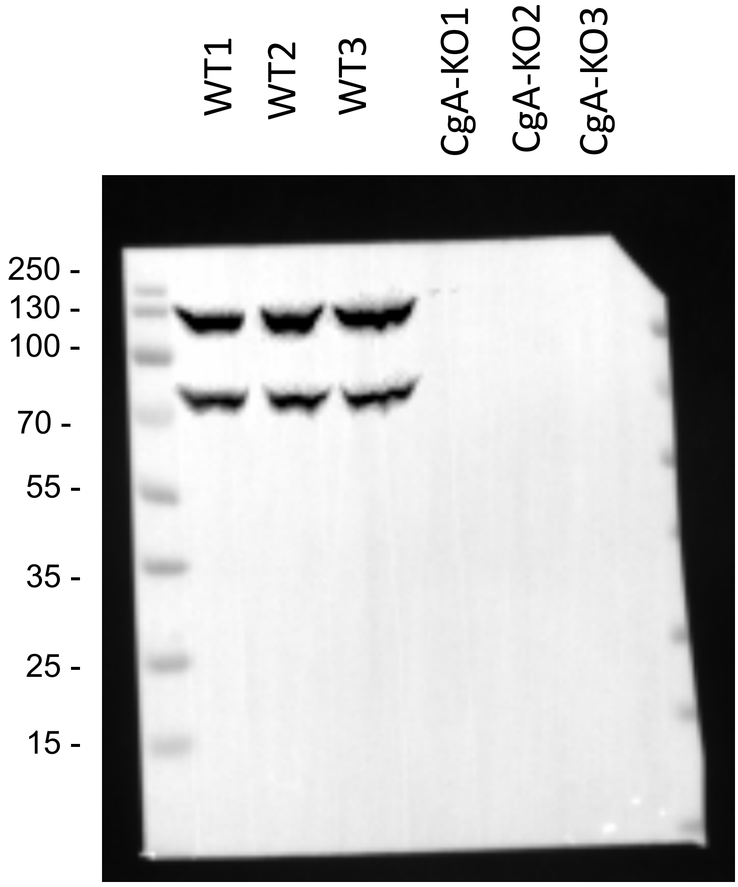


CgA staining

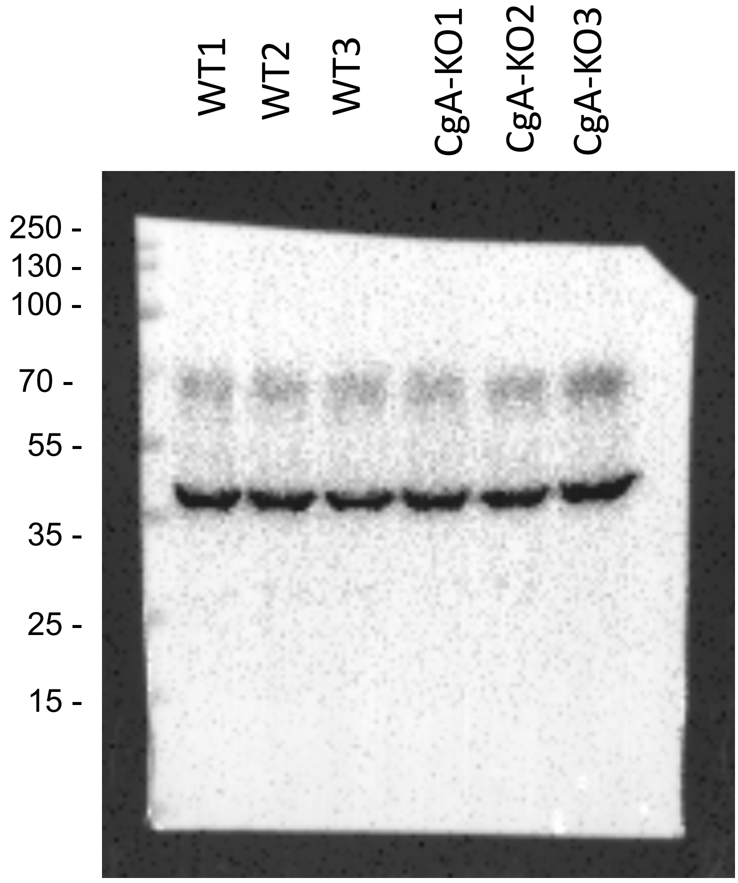


Actin staining

**S-Fig.6 Original immunoblots from sup Fig. 4**
Original immunoblotting images showing chromogranin A or actin staining in WT and CgA-KO mice.

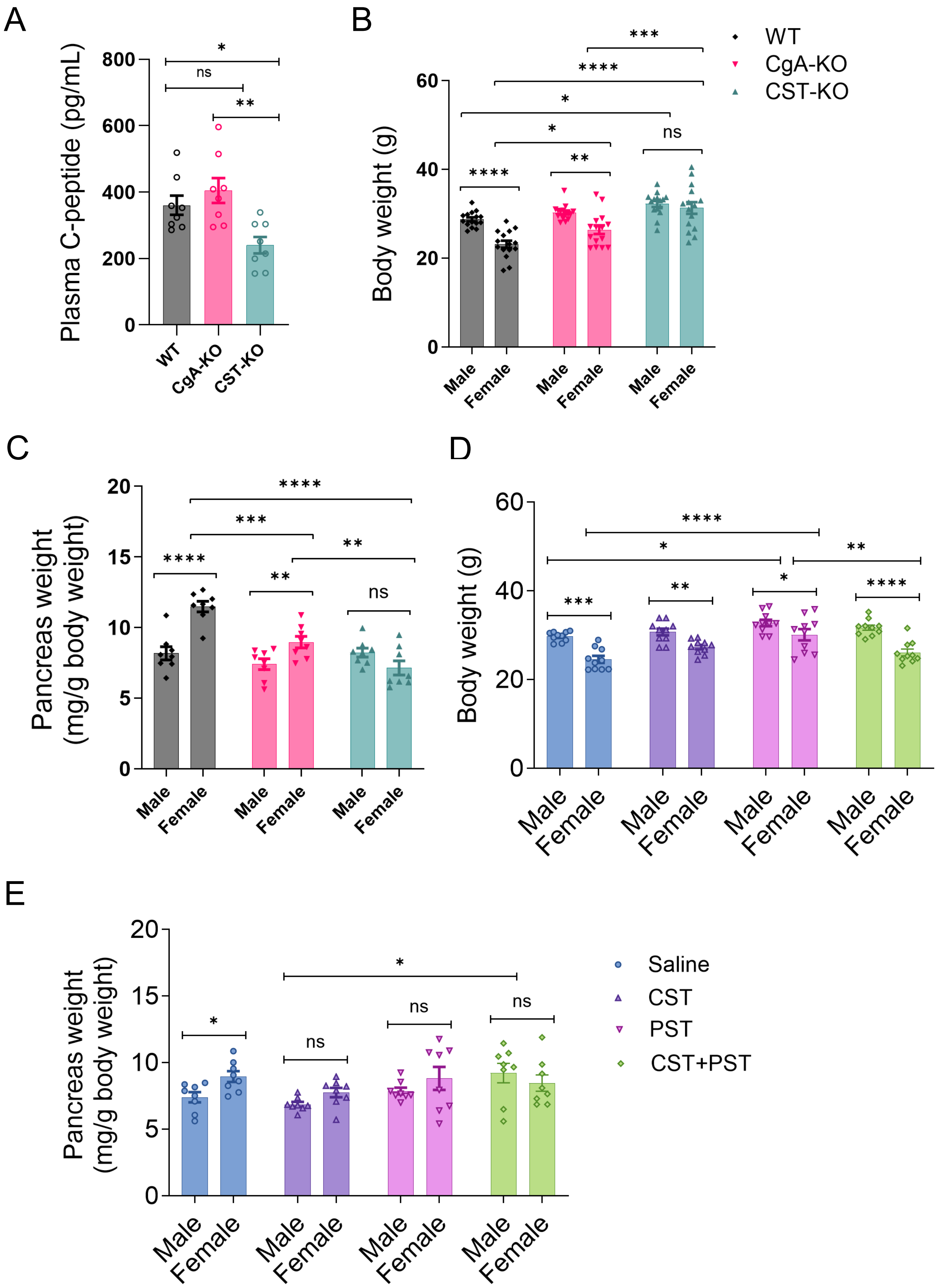


**S-Fig.7 Mice characteristics: peptide levels, pancreatic weight and body weight**
**A)**.C-peptide plasma levels (pg/mL). **B)** Male and female body weight (g) for WT, CgA-KO or CST-KO mice. N=15 **C) )** Male and female pancreatic weight (mg/g body weight) for WT, CgA-KO or CST-KO mice. N=15 **D)** Body or pancreatic **E)** weight of CgA-KO male and female mice supplemented with saline, CST, PST or CST+PST. N=10. *p<0.05 , **p<0.01, ***p<0.001, ****p<0.0001

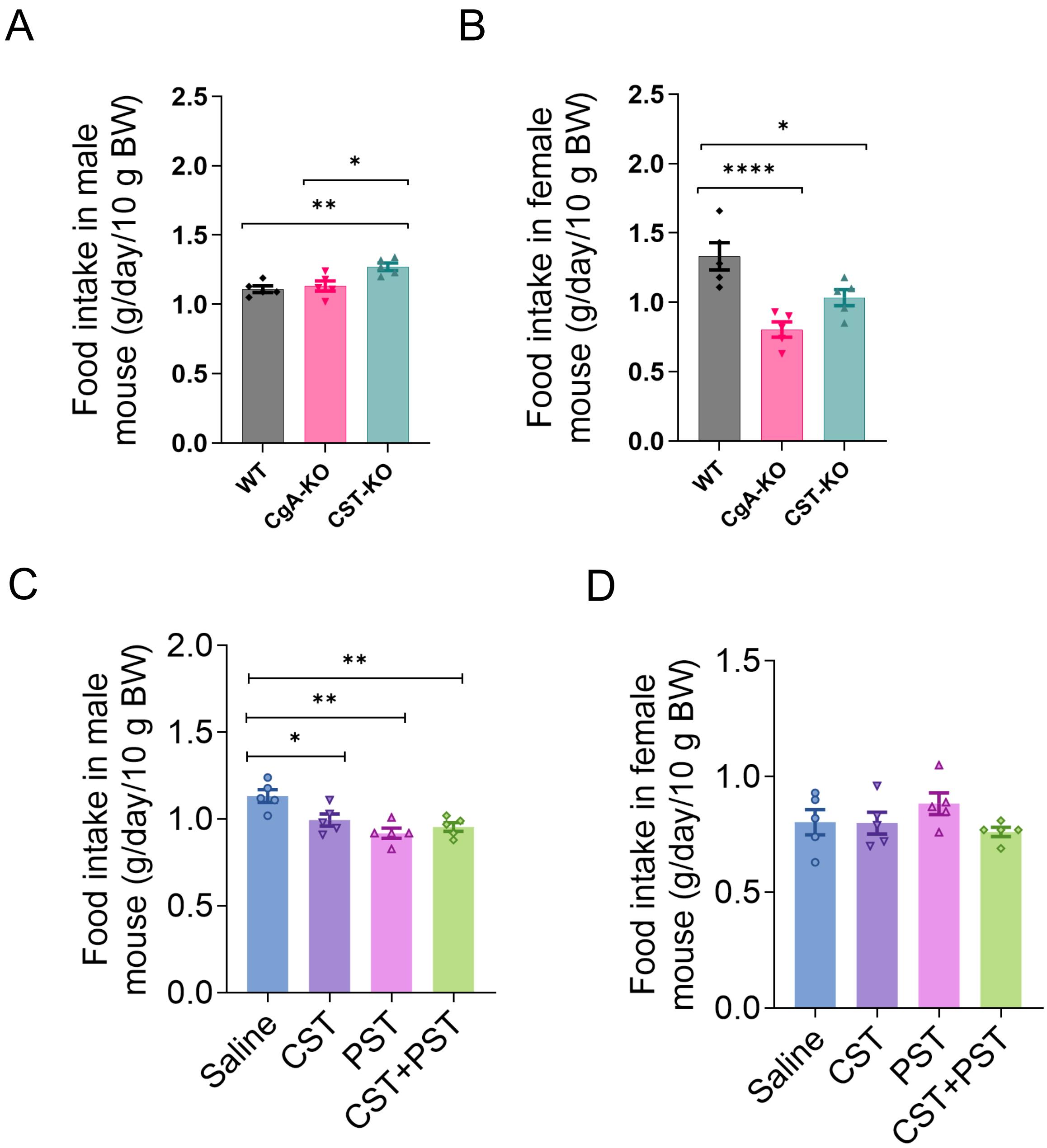


**S-Fig.8 Mice characteristics: food intake**
**A-B)**.Male and female food intake (g/day/10 g BW) measured for WT, CgA-KO or CST-KO mice. N=5 **C-D) )** Male and female food intake (g/day/10 g BW) measured for CgA-KO male and female mice supplemented with saline, CST, PST or CST+PST. N=5. *$p<0.05$ , **$p<0.01$, ***$p<0.001$

A

| Time (min) | WT vs CgA-KO | WT vs CST-KO | CgA-KO vs CST-KO |
|---|---|---|---|
| 0 | ns | ns | ns |
| 15 | * | ** | **** |
| 30 | *** | ** | **** |
| 60 | ** | ** | **** |
| 90 | ns | * | *** |
| 120 | ns | ns | ** |

B

| Time (min) | WT vs CgA-KO | WT vs CST-KO | CgA-KO vs CST-KO |
|---|---|---|---|
| 0 | ns | ns | ns |
| 15 | ns | ** | **** |
| 30 | ** | *** | **** |
| 60 | *** | *** | **** |
| 90 | ** | *** | **** |
| 120 | * | ** | **** |

C

| | Significance of CgA-KO supplemented with | | | | | |
|---|---|---|---|---|---|---|
| Time (min) | Sal vs CST | Sal vs PST | Sal vs CST+PST | CST vs PST | CST vs CST+PST | PST vs CST+PST |
| 0 | ns | ns | ns | ns | ns | ns |
| 15 | ns | ** | ns | **** | ns | ** |
| 30 | ns | **** | ns | **** | * | **** |
| 60 | ns | **** | ns | **** | ns | **** |
| 90 | ns | **** | ns | **** | ns | **** |
| 120 | ns | *** | ns | **** | ns | *** |

**S-Fig.9 Significance Male GTT and PTT data from main Fig. 3**
**A)** GTT significance table from GTT graph panel A comparing blood glucose levels (mg/dL) over time (min) of WT, CgA-KO and CST-KO male mice. **B)** Pyruvate tolerance test (PTT) significance table from PTT graph panel C comparing blood glucose levels of WT, CgA-KO and CST-KO male mice. **C)** PTT significance table from PTT graph showing blood glucose results (mg/dL) over time (min) of male CgA-KO mice supplemented with saline, CST, PST or CST+PST N=10 per group. *$p<0.05$ , **$p<0.01$, ***$p<0.001$, ****$p<0.0001$

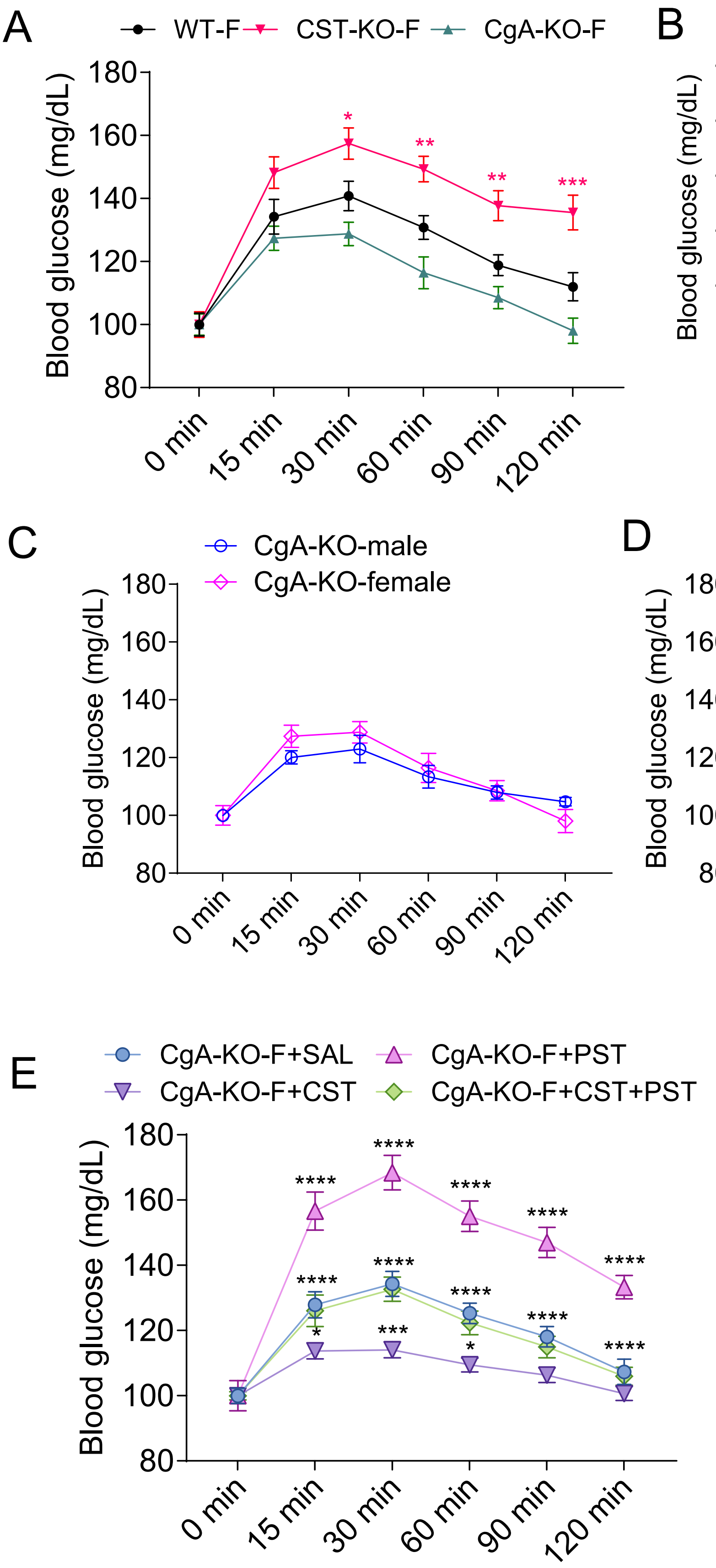


**S-Fig.10 Female PTT data and male and female comparison**
**A)** Graph displaying pyruvate tolerance test (PTT) blood glucose results (mg/dL) for female (F) WT (black), CgA-KO (pink) and CST-KO (green) over time (min) N=15 per group. **B-D)** Graphs displaying combined data of PTT, comparing males and females WT, CgA-KO and CST-KO blood glucose results (mg/dL) over time (min). **E)** Graph displaying combined data of PTT blood glucose results (mg/dL) over time (min) of female (F) CgA-KO mice supplemented with saline, CST, PST or CST+PST N=10 per group. *p<0.05, **p<0.01, ***p<0.001, ****p<0.0001

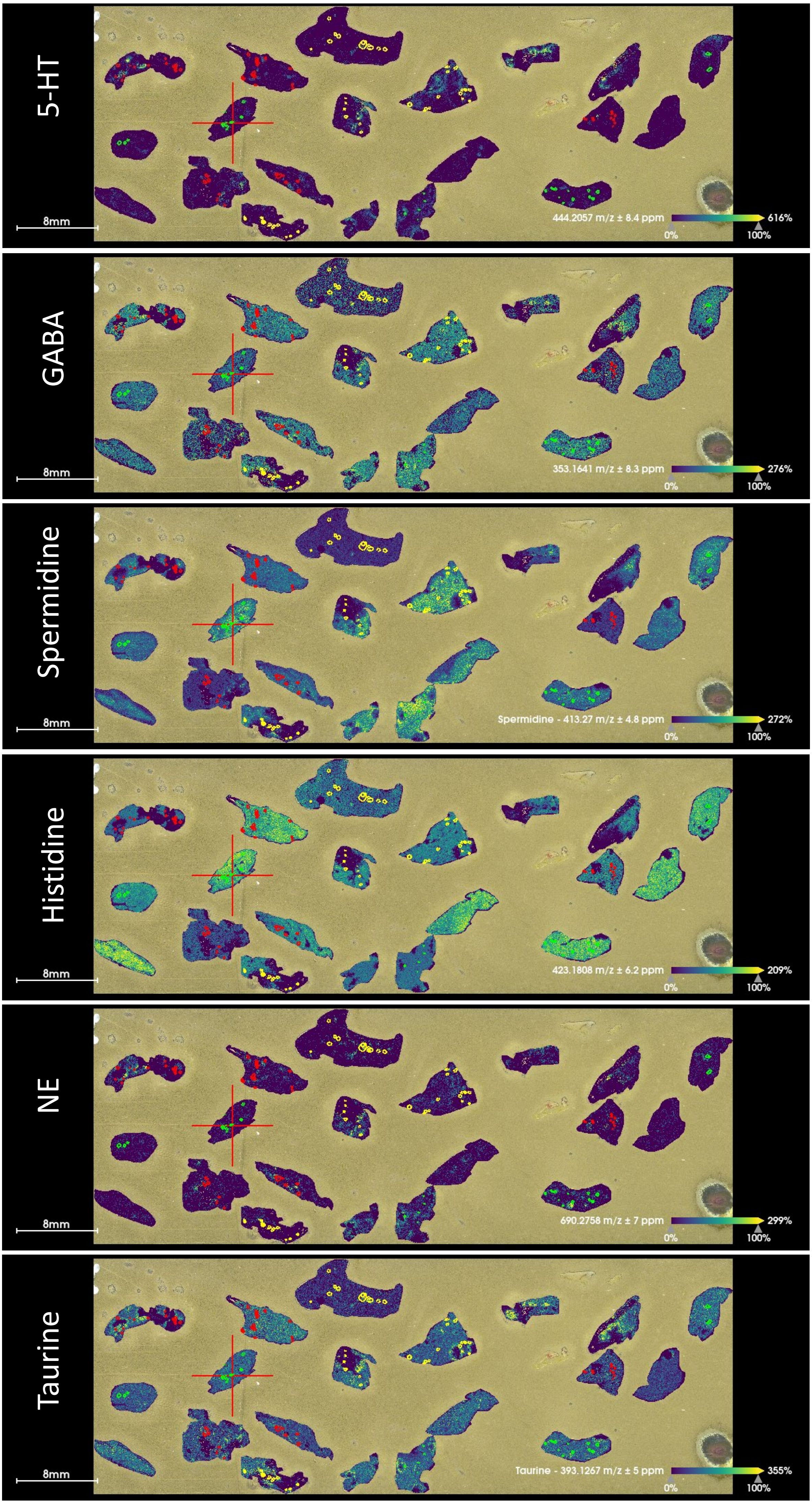


**S-Fig.11 Spatial MS heatmaps of analytes identified in the pancreas**
Spatial MS full heatmaps showing all measured pancreata including annotated islet and exocrine areas for WT (red), CgA-KO (green), CST-KO (yellow).

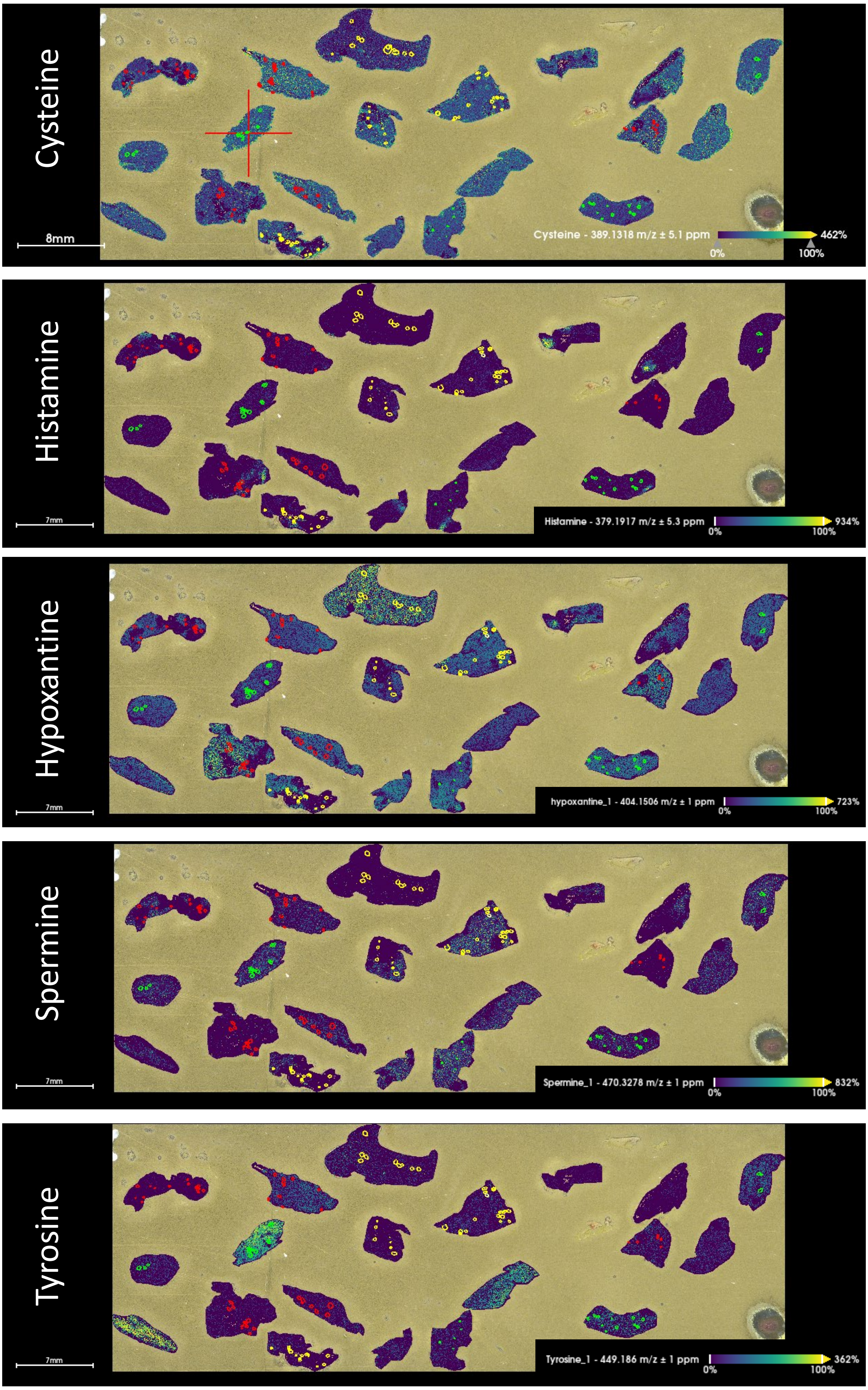


**S-Fig. 12 Spatial MS heatmaps of analytes identified in the pancreas**
Spatial MS full heatmaps showing all measured pancreata including annotated islet and exocrine areas for WT (red), CgA-KO (green), CST-KO (yellow).

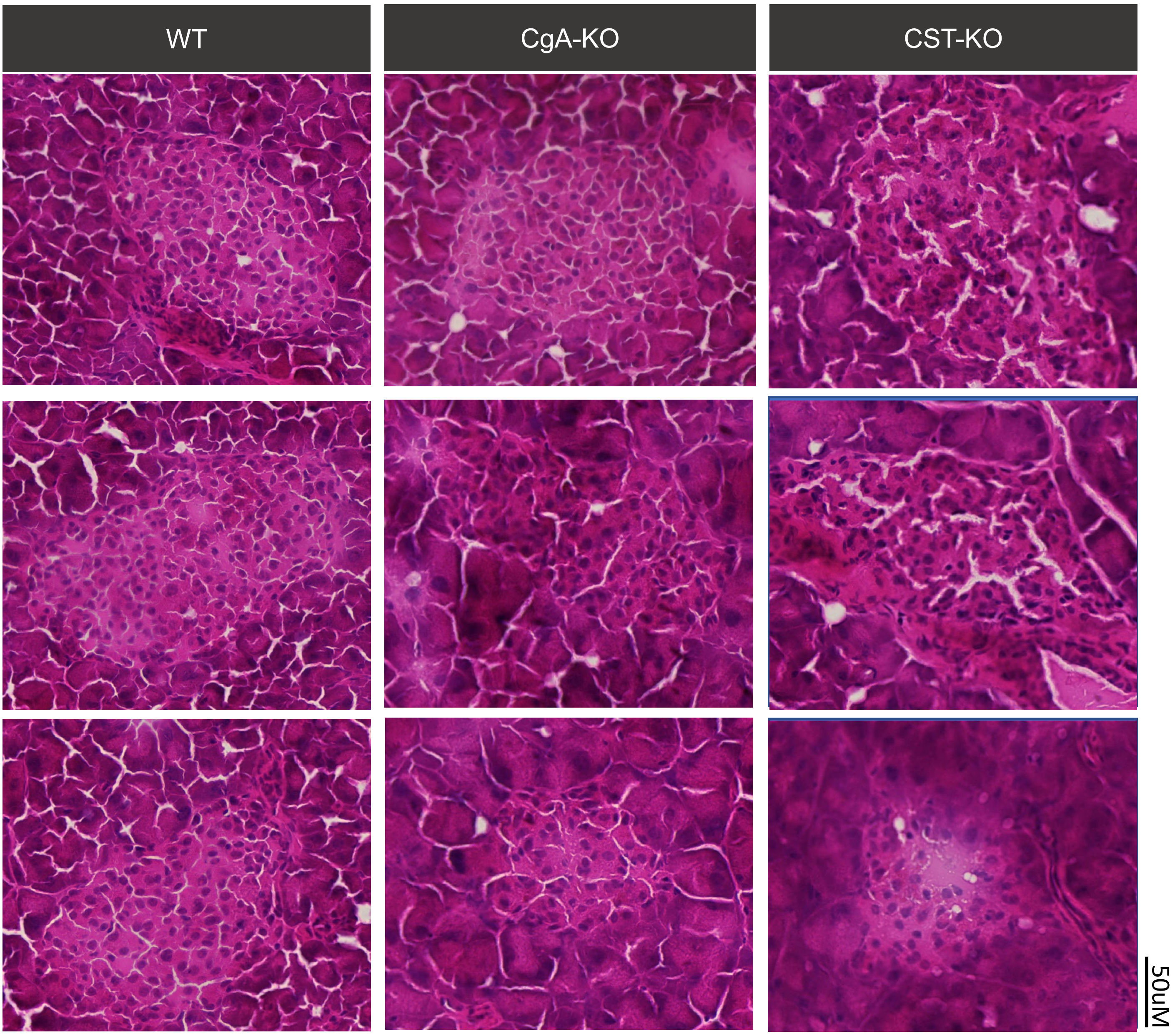


**S-Fig. 13 H&E stain for spatial MS of male pancreatic islets.**
Consecutive sections from these were used for mass spectrometry imaging and the H&E stainings here guided islet annotations (see S-Method Fig 2)

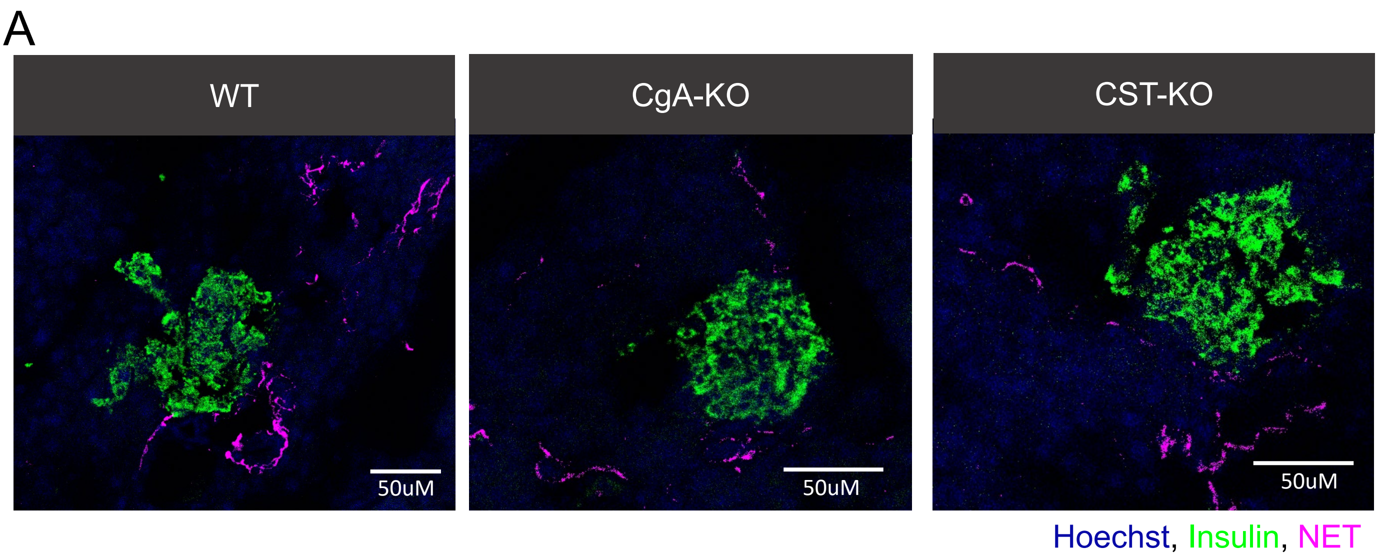


**S-Fig.14 Female innervation of pancreatic islets and plasma cytokines**
**A)** Representative images of nerve staining (norepinephrine transporter (NET, magenta), islets (insulin, green) and nuclei (Hoechst) in WT, CgA-KO and CST-KO pancreatic slices.

### Annotation of islets based on staining in all channels

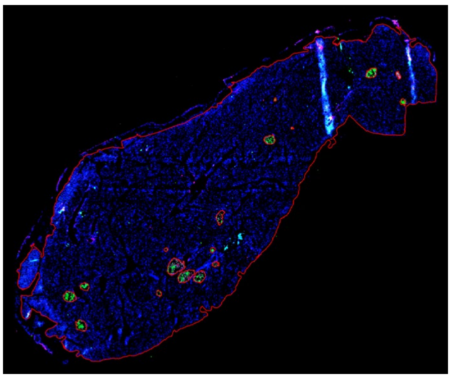

### Perform cell detection on annotated islets by training classifier

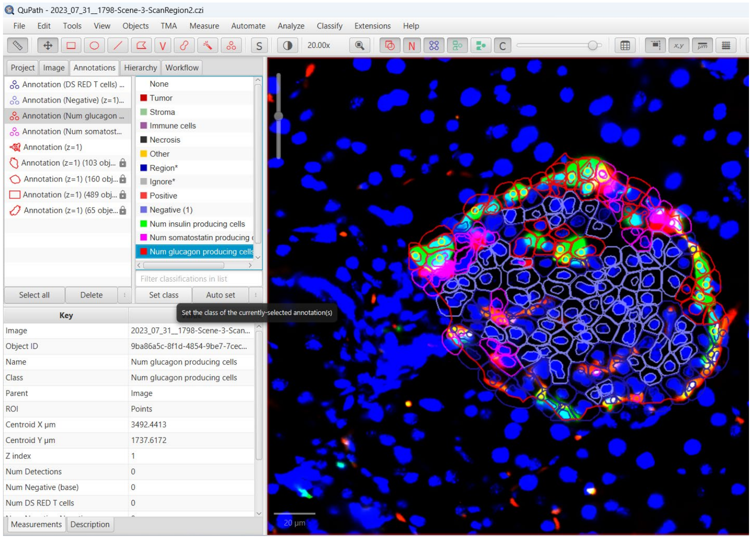

Settings for cell detection

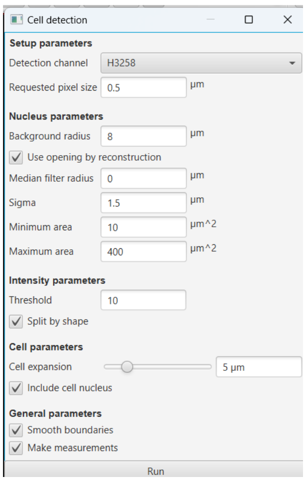


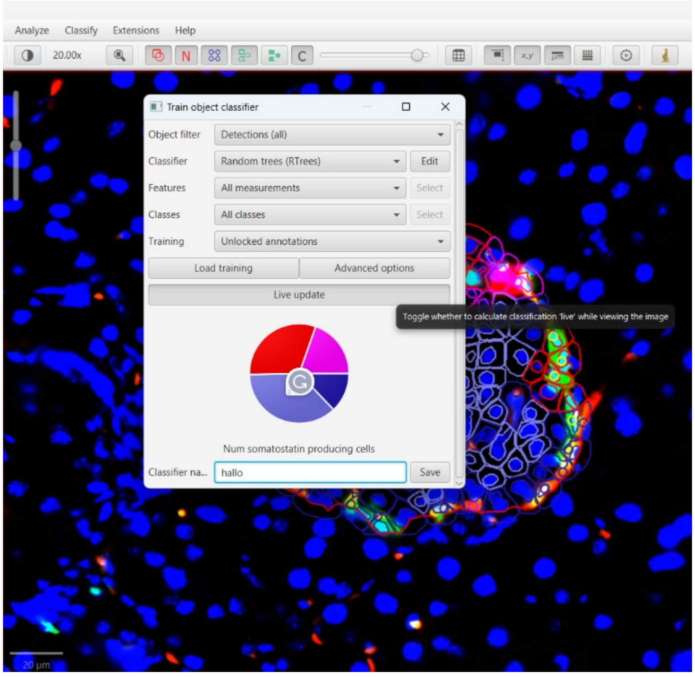


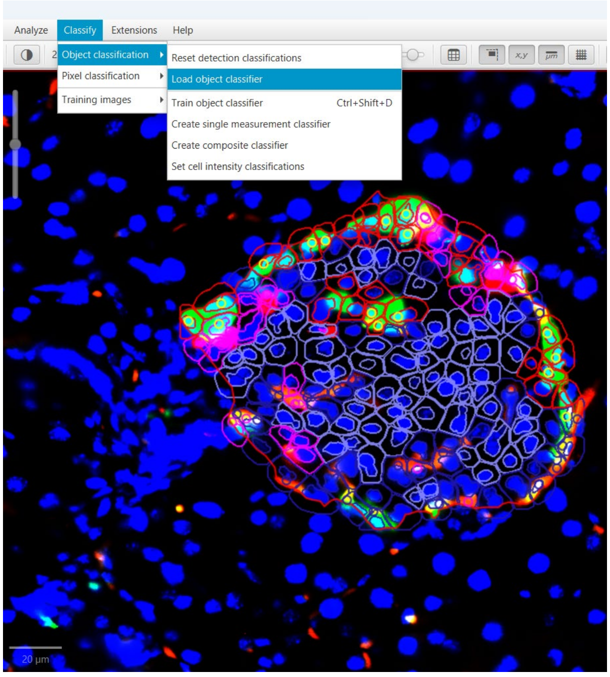


### Example of classified islet

Obtained cell detection using classifier

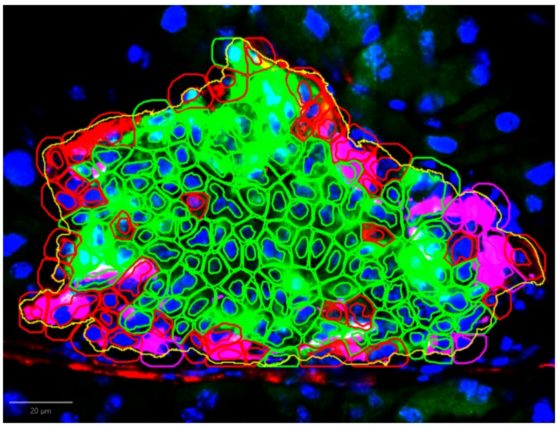

Legend
Insulin
Glucagon
Somatostatin
Negative

Original fluorescent staining

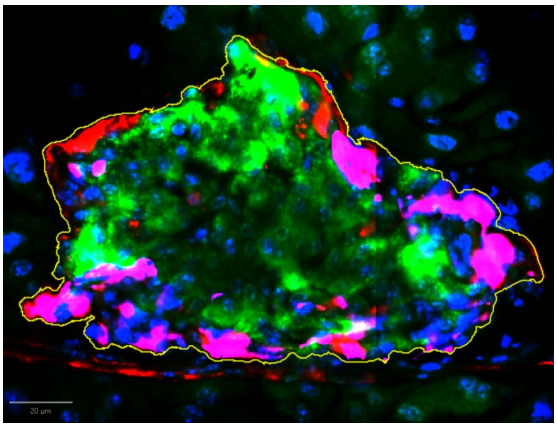

**S-Method Fig.1 QuPath workflow for alpha/beta/delta cell detection**
Step-by-step workflow for cell detection, annotation and quantification.

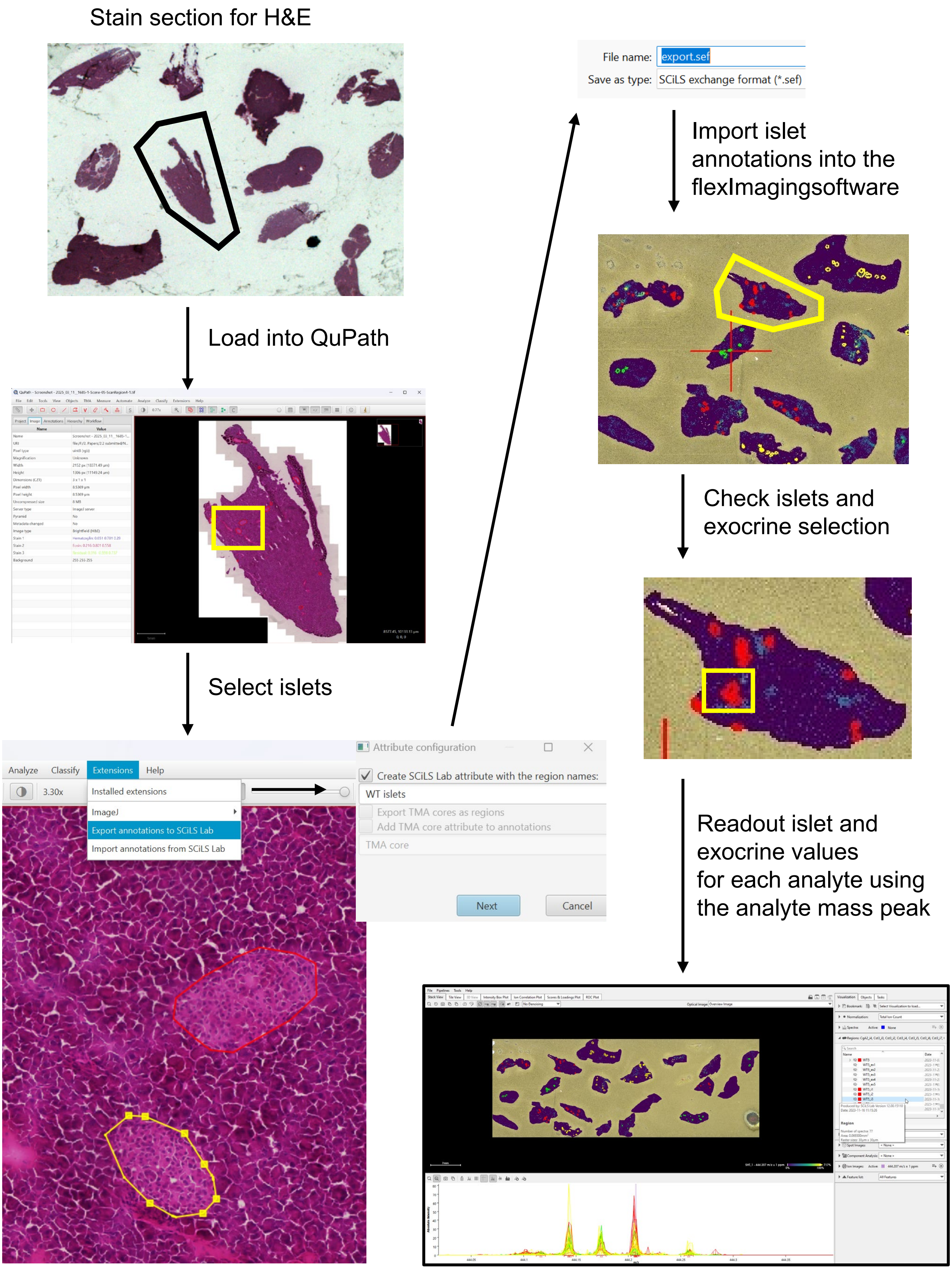


**S-Method Fig. 2 Spatial MS workflow**
Step-by-step workflow for islet annotation in spatial MS data using both H&E stained consecutive sections and serotonin quantity.

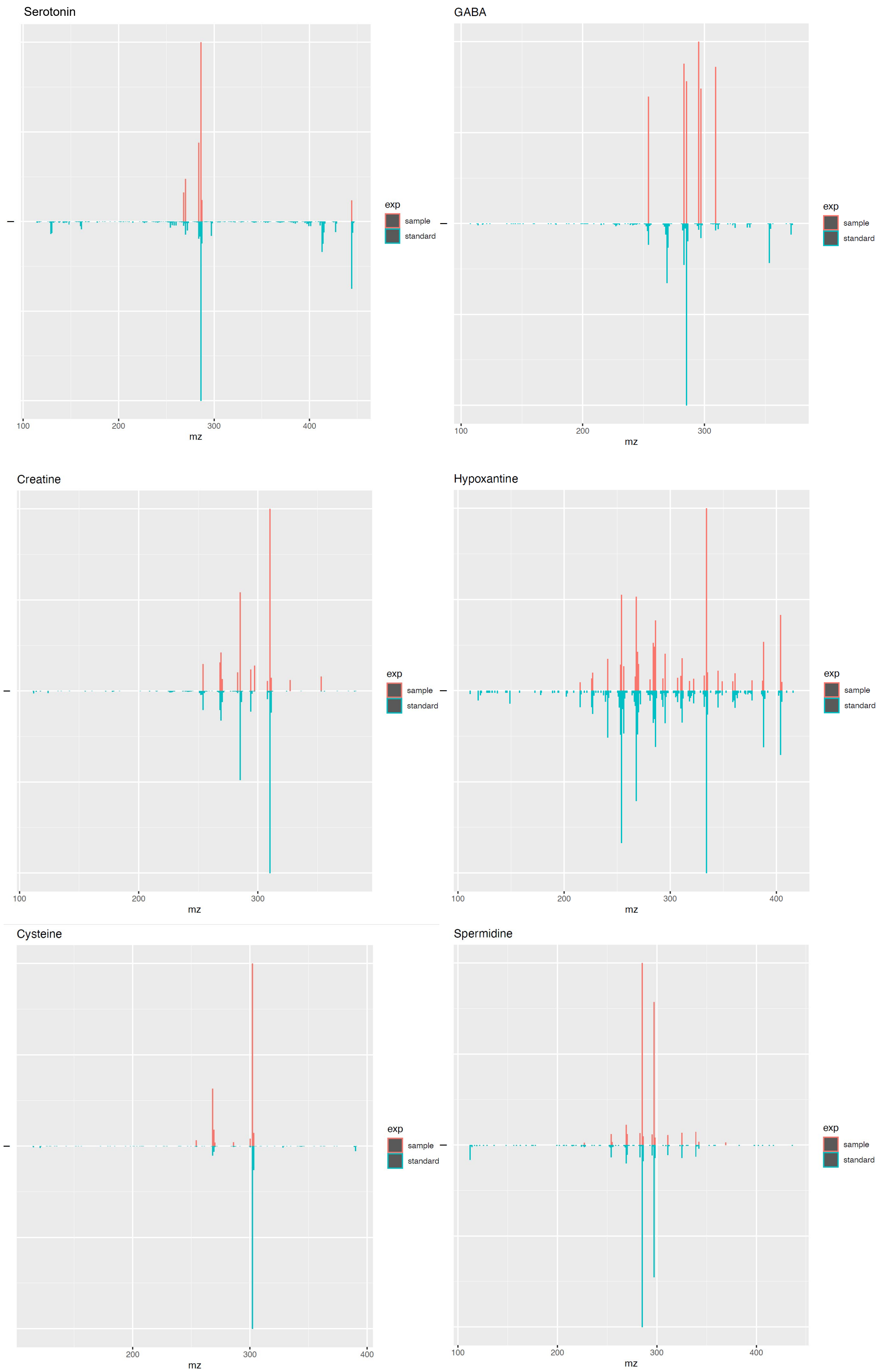


**S-Method Fig. 3 MS/MS plots of analytes identified in the pancreas**
Red lines show pancreas sample signal and blue lines show signal of the corresponding control standard.

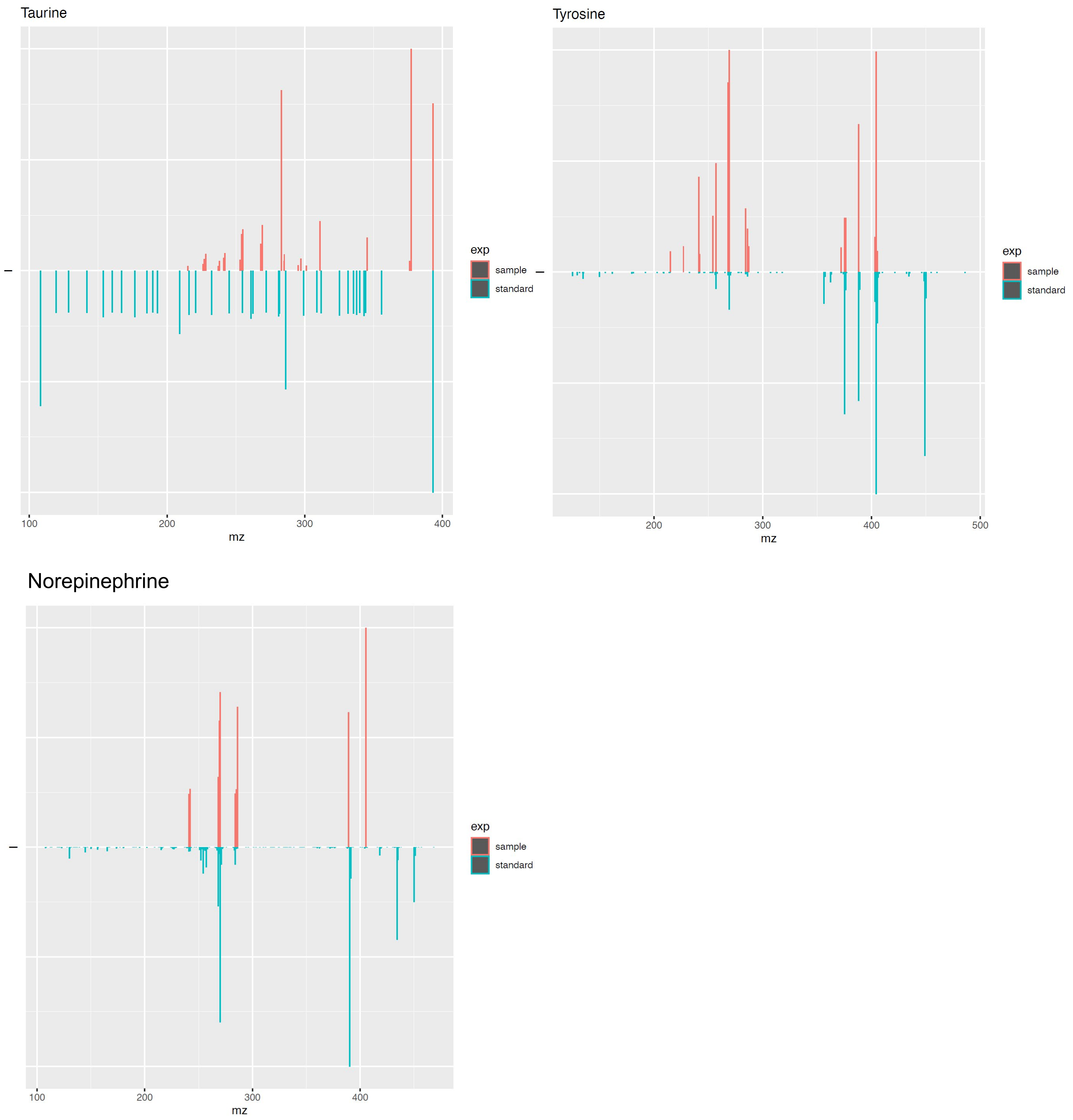


**S-Method Fig. 4 MS/MS plots of analytes identified in the pancreas 2**
Red lines show pancreas sample signal and blue lines show signal of the corresponding control standard.
